# A young age of subspecific divergence in the desert locust *Schistocerca gregaria*, inferred by ABC Random Forest

**DOI:** 10.1101/671867

**Authors:** Marie-Pierre Chapuis, Louis Raynal, Christophe Plantamp, Christine N. Meynard, Laurence Blondin, Jean-Michel Marin, Arnaud Estoup

## Abstract

Dating population divergence within species from molecular data and relating such dating to climatic and biogeographic changes is not trivial. Yet it can help formulating evolutionary hypotheses regarding local adaptation and future responses to changing environments. Key issues include statistical selection of a demographic and historical scenario among a set of possible scenarios, and estimation of the parameter(s) of interest under the chosen scenario. Such inferences greatly benefit from new statistical approaches including approximate Bayesian computation - Random Forest (ABC-RF), the latter providing reliable inference at a low computational cost, with the possibility to take into account prior knowledge on both biogeographical history and genetic markers. Here, we used ABC-RF, including independent information on evolutionary rate and pattern at microsatellite markers, to decipher the evolutionary history of the African arid-adapted pest locust, *Schistocerca gregaria*. We found that the evolutionary processes that have shaped the present geographical distribution of the species in two disjoint northern and southern regions of Africa were recent, dating back 2.6 Ky (90% CI: 0.9 – 6.6 Ky). ABC-RF inferences also supported a southern colonization of Africa from a low number of founders of northern origin. The inferred divergence history is better explained by the peculiar biology of *S. gregaria*, which involves a density-dependent swarming phase with some exceptional spectacular migrations, rather than a continuous colonization resulting from the continental expansion of open vegetation habitats during more ancient Quaternary glacial climatic episodes.

## Introduction

As in other regions of the world, Africa has gone through several major episodes of climate change since the early Pleistocene (deMenocal 1995 and 2004). The prevalent climate was colder and drier than nowadays during glaciation periods, and became more humid during warmer interglacial periods. These climatic phases resulted in shifts of vegetation (de Vivo and Carmignotto 2004) and are most likely at the origin of the current isolation between northern and southern distributions of arid-adapted species (Monod 1971). In Africa, at least fifty-six plant species show disjoint geographical distributions in southern and northern arid areas (Monod 1971; Jurgens 1997; Lebrun 2001). Similarly, a number of animal vertebrate species show meridian disjoint distributions on this continent, including eight mammals and 29 birds (Monod 1971; de Vivo and Carmignotto 2004; Lorenzen *et al*. 2012). The desert locust, *Schistocerca gregaria*, is among the few examples of insect species distributed in two distinct regions along the north-south axis of Africa. Other known disjunctions in insects are interspecific and concern species of the families Charilaidae (Orthoptera) and Mythicomyiidae (Diptera), and of the genus *Fidelia* (Hymenoptera) (Le Gall *et al*. 2010). Similarities in extant distributions of African arid-adapted species across divergent taxonomic groups point to a common climatic history and an important role of environmental factors. Yet, to our knowledge, studies relating evolutionary history and climatic history have rarely been carried out in this continent; but see mitochondrial studies by Miller *et al*. 2011 on the ostrich, Atickem *et al*. 2018 on the black-backed jackal, and Moodley *et al*. 2018 on the white rhinoceros.

Relating evolutionary and climatic histories often requires dating population differentiation events so that species or subspecies divergence can be understood in a broad biogeographic context. However, finding a reliable calibration to convert measures of genetic divergence into units of absolute time is challenging, especially so for recent evolutionary events (Ho *et al*. 2008). Internal fossil records are often lacking and extra-specific fossil calibration may lead to considerable overestimates of divergence times (Ho *et al*. 2008). A sensible approach is to use an evolutionary rate estimated from sequence data of a related species for which internal fossil calibration is available (Ho *et al*. 2008). Unfortunately, on the African continent, fossils, such as radiocarbon-dated ancient samples, remain relatively rare and are often not representative of modern lineages (*e*.*g*., Le Gall *et al*. 2010 for insects). The lack of paleontological and archaeological records is partly due to their fragility under the aridity conditions of the Sahara. The end-result is that the options to relate population divergence to biogeographic events in this region are very limited. Finding a dating strategy that does not rely on fossils would therefore constitute a key advance in understanding the region’s biogeography.

In this context, the use of versatile molecular markers, such as microsatellite loci, for which evolutionary rates can be obtained from direct observation of germline mutations in the species of interest, represents a useful alternative. Microsatellite mutation rates exceed by several orders of magnitude that of point mutation in DNA sequences, ranging from 10^−6^ to 10^−2^ events per locus and per generation (Ellegren 2000). Such rates allow one to both observe mutation events in parent-offspring segregation data of realistic sample size and to reconstruct the recent history of related populations. However, the use of microsatellite loci to estimate divergence times at recent evolutionary time-scales still needs to overcome significant challenges. Since microsatellite allele sizes result from the insertion or deletion of single or multiple repeat units and are tightly constrained, these markers can be characterized by high levels of homoplasy that can obscure inferences about gene history (*e*.*g*., Estoup *et al*. 2002). In particular, at large time scales (*i*.*e*., for distantly related populations), genetic distance values no longer follow a linear relationship with time. Rather, they reach a plateau and therefore provide biased and hence unreliable estimation of divergence time over a certain time threshold (Takezaki and Nei 1996; Feldman *et al*. 1997; Pollock *et al*. 1998). Microsatellites remain informative with respect to divergence time only if the population split occurs within the period of linearity with time (Feldman *et al*. 1997; Pollock *et al*. 1998). The exact value of the differentiation threshold above which microsatellite markers would no longer accurately reflect divergence times will depend on constraints on allele sizes and population-scaled mutation rates (Feldman *et al*. 1997; Pollock *et al*. 1998). In this context, the approximate Bayesian computation - random forest (ABC-RF) approach recently proposed by Raynal *et al*. (2019) is a singular statistical advance, as it allows both to take into account prior knowledge on genetic markers (including mutation rate and pattern) and to compute accuracy of parameter estimation at a local (*i*.*e*., posterior) scale. Using this methodological framework, one can envisage to evaluate the divergence time threshold above which posterior estimates of divergence time would become biased under a given evolutionary scenario, and hence in this way thoroughly evaluate the robustness of inferences about divergence time events.

The desert locust, *S. gregaria*, is a generalist herbivore that can be found in arid grasslands and deserts in both northern and southern Africa (Figure 1a). In its northern range, the desert locust is one of the most widespread and harmful agricultural pest species, with a very large potential outbreak area, spanning from West Africa to Southwest Asia. The desert locust is also present in the south-western arid zone (SWA) of Africa, which includes South-Africa, Namibia, Botswana and south-western Angola. The southern populations of the desert locust are termed *S. g. flaviventris* and are geographically separated by nearly 2,500 km from populations of the nominal subspecies from northern Africa, *S. g. gregaria* (Uvarov 1977). The isolation of *S. g. flaviventris* and *S. g. gregaria* lineages was recently supported by highlighting distinctive mitochondrial DNA haplotypes and male genitalia morphologies (Chapuis *et al*. 2016). Yet, the precise history of divergence remains elusive.

**Figure 1.**
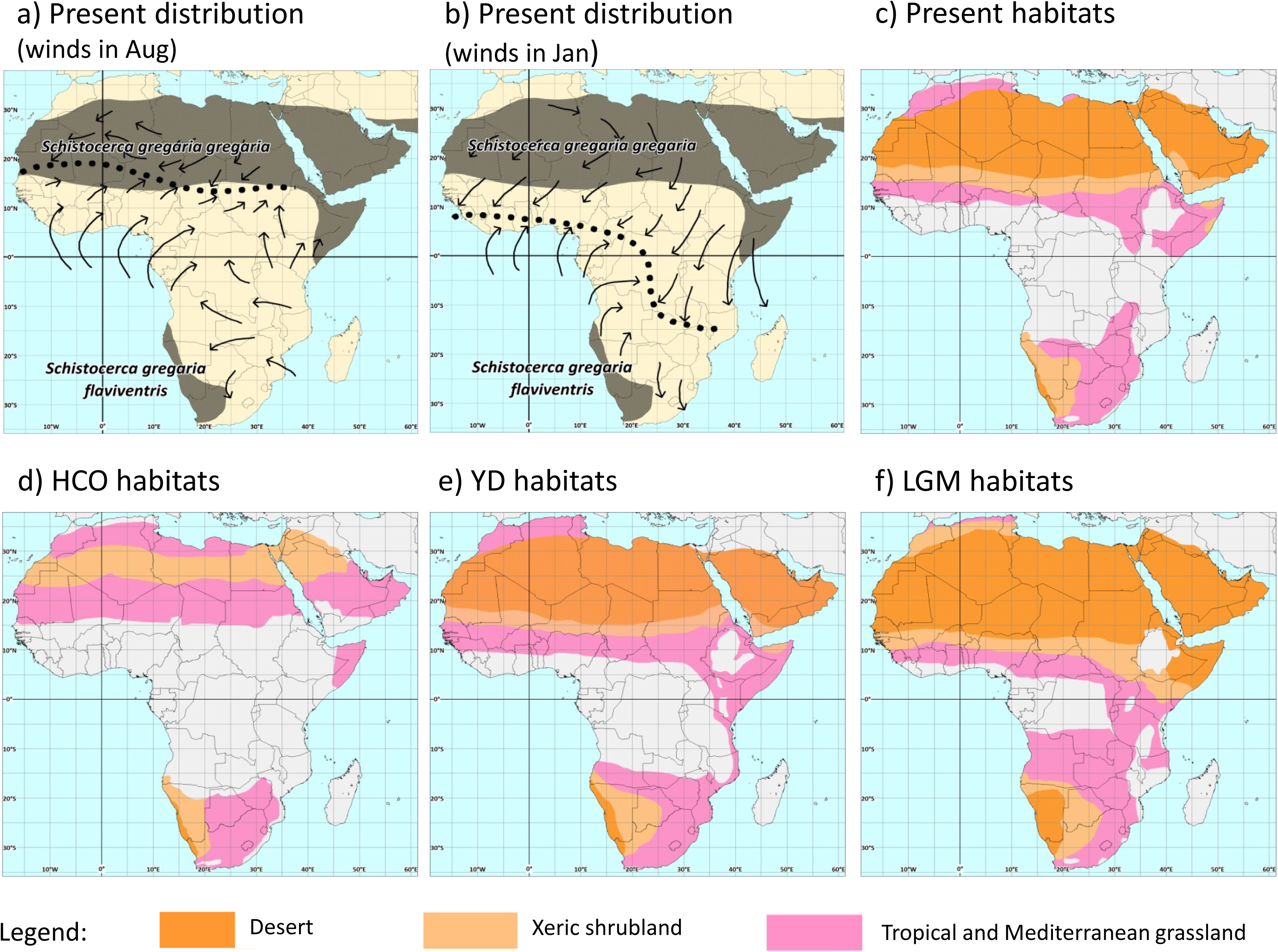
Present time distribution range of Schistocerca gregaria in Africa under remission periods with winds in August a) and January b), and vegetation habitats suitable for the species during the present period c), the Holocene Climatic Optimum (HCO, 9 to 6 Ky ago) d), the Younger Dryas (YD, 12.9 to 11.7 Ky ago) e) and the Last Glacial Maximum (LGM, 26 to 14.8 Ky ago) f). (a-b) Distribution range and winds are adapted from Sword et al. (2010) and Nicholson (1996), respectively. (c-f) Vegetation habitats are adapted from Adams and Faure (1997). Open vegetation habitats suitable for the desert locust correspond to deserts (dark orange), xeric shrublands (light orange) and tropical - Mediterranean grasslands (pink). Other unsuitable habitat classes (white) are forests, woodlands and temperate shrublands and savannas.

The main objective of the present study is to unravel the historical and evolutionary processes that have shaped the present disjoint geographical distribution of the desert locust and the genetic variation observed both within and between populations of its two subspecies. To this aim, we applied an ABC-RF approach and show its full potential to help discriminate between alternative biogeographic scenarios. We first start by identifying a set of evolutionary alternatives relevant to the species from African paleo-vegetation maps that reflect potential past distributions of the desert locust. We then used molecular data obtained from microsatellite markers for which we could obtain independent information on evolutionary rates and allele size constraints in the species of interest from direct observation of germline mutations (Chapuis *et al*. 2015). We applied recently available algorithms of the ABC-RF on our microsatellite population genetic data to compare a set of thoroughly formalized and justified evolutionary scenarios and estimate the divergence time between *S. g. gregaria* and *S. g. flaviventris* under the most likely of our scenarios. Finally, we interpret our results in light of past vegetation cover and desert locust biology.

## Results

Table 1 shows the values of the summary statistics obtained from the observed population dataset consisting in two unstructured pooled samples of the subspecies *S. g. gregaria* and *S. g. flaviventris*. A total of 170 individuals (*i*.*e*., 80 and 90 individuals for *S. g. gregaria* and *S. g. flaviventris*, respectively) were genotyped at 23 microsatellite markers derived from either genomic DNA (14 loci) or messenger RNA (9 loci) resources (hereafter referred to as untranscribed and transcribed microsatellite markers, respectively). The level of differentiation between the two subspecies (as measured by the parameter *F*_ST_) was 0.04 and 0.12 for untranscribed and transcribed microsatellite markers, respectively. The level of genetic diversity was higher within the northern subspecies *S. g. gregaria* (+7% and +14% for the mean number of alleles and expected heterozygosity, respectively).

**Table 1.**
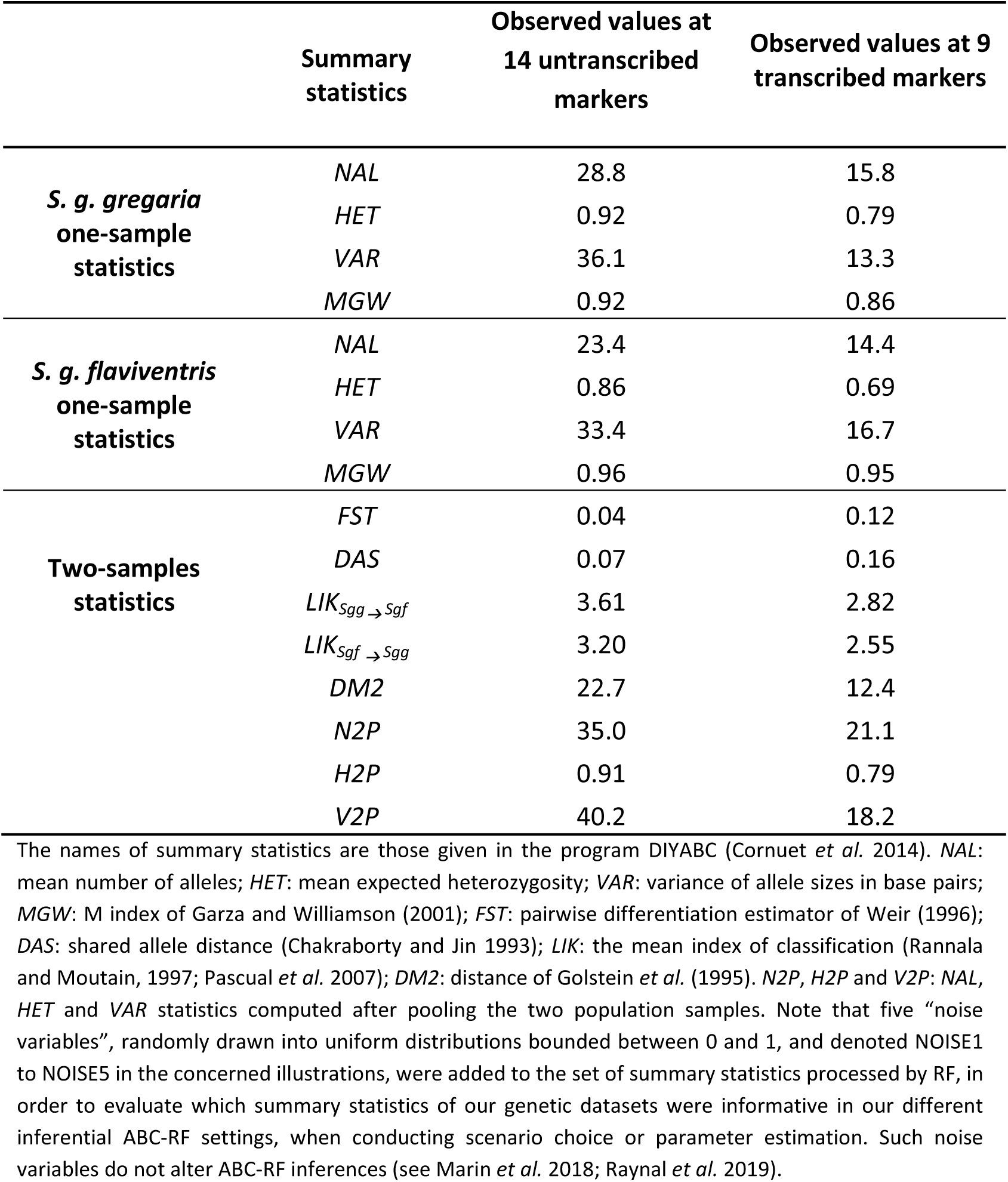
Summary statistics provided by DIYABC with corresponding values computed from the observed microsatellite dataset.

### Formalization and choice of evolutionary scenarios

Using a rich corpus of vegetation data, we reconstructed the present time (Fig. 1c) and past time (Figs. 1d-f) distribution ranges of *S. gregaria* in Africa, going back to the Last Glacial Maximum period (LGM, 26 to 14.8 Ky ago). Maps of vegetation cover for glacial arid maximums (Figs. 1e and 1f) showed an expansion of open vegetation habitats sufficient to make the potential range of the species continuous from the Horn of Africa in the north-west to the Cape of Good Hope in the south. Maps of vegetation cover for interglacial humid maximums (Fig. 1d) showed a severe contraction of deserts. These maps helped us formalize twelve competing evolutionary scenarios (Figure 2), as well as bounds of prior distributions for various parameters (see the section *Prior setting for historical and demographical parameters* in Materials and methods). The twelve competing scenarios were generated by the combinations of presence *vs*. absence of three key evolutionary events that we identified as having potentially played a role in setting up the observed disjoint distribution of the two locust subspecies: (i) a long population size contraction in the ancestral population, due to the reduction of open vegetation habitats during the interglacial periods, (ii) a bottleneck in the southern subspecies *S. g. flaviventris* right after divergence reflecting a single long-distance migration event of a small fraction of the ancestral population, and (iii) a discrete genetic admixture event either unidirectional from the ancestral northern subspecies *S. g. gregaria* into *S. g. flaviventris*, or bidirectional, in order to consider the many climatic transitions of the last Quaternary.

**Figure 2.**
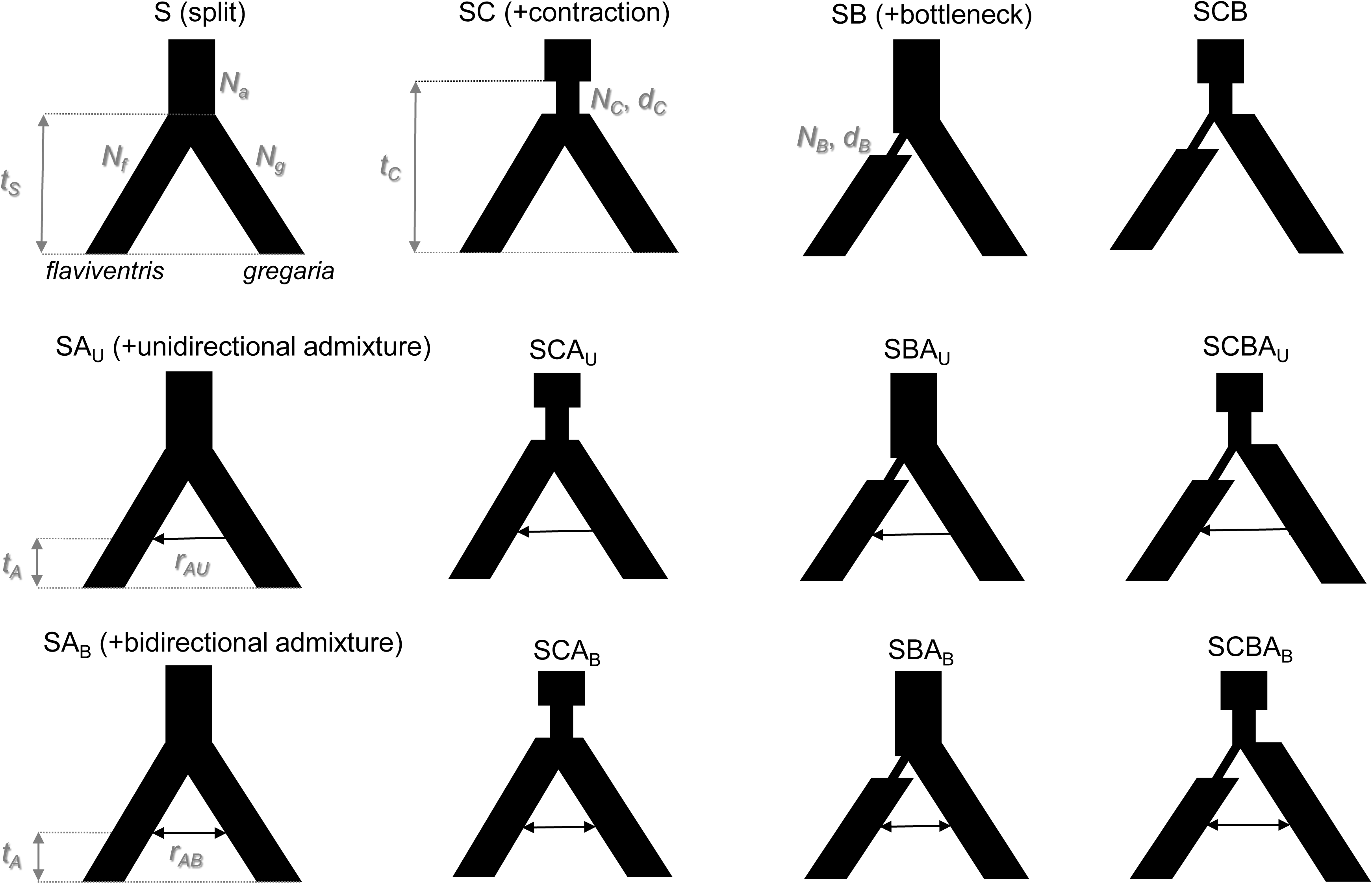
Evolutionary scenarios compared using ABC-RF. The subscripts g, f and a refer to the subspecies S. g. gregaria, S. g. flaviventris and their unsampled common ancestor, respectively. Twelve scenarios are considered and identified by an acronym. Such scenarios differ by the presence or absence of three evolutionary events: a bottleneck in S. g. flaviventris (B) right after divergence between the two subspecies, a population size contraction in the ancestral population (C) and a discrete genetic admixture event either unidirectional from S. g. gregaria into S. g. flaviventris (AU) or bidirectional (AB). For convenience, parameters associated with an evolutionary event are represented graphically only in the first shown scenario that include this event. Looking forward in time, time periods are tC, the time of ancestral population size contraction, tS, the time of split between the two subspecies, and tA, the time of the genetic admixture between subspecies (with tC > tS > tA). rAU is the unidirectional admixture rate, i.e. the proportion of genes from the S. g. gregaria lineage entering the S. g. flaviventris population at time tA. rAB is the bidirectional admixture rate, i.e. the proportion of genes exchanged between the S. g. gregaria lineage and the S. g. flaviventris lineage at time tA. Ng, Nf and Na are the stable effective population sizes of S. g. gregaria, S. g. flaviventris and the ancestor, respectively. NC is the effective population size during the contraction event of duration dC in the ancestor. NB is the effective population size during the bottleneck event of duration dB.

ABC-RF analyses supported the same group of scenarios or the same best individual scenario for all ten replicate analyses (Table 2). The classification votes and posterior probabilities estimated for the observed microsatellite dataset were the highest for the groups of scenarios in which (i) *S. g. flaviventris* experienced a bottleneck event at the time of the split (scenarios SB+SCB+SBA_U_+SBA_B_+SCBA_U_+SCBA_B_ in Figure 2; average of 2,829 votes out of 3,000 RF-trees; posterior probability = 0.926), (ii) the ancestral population experienced a population size contraction (scenarios SC+SCB+SCA_U_+SCA_B_+SCBA_U_+SCBA_B_; 2,035 of 3,000 RF-trees; posterior probability = 0.682), and (iii) no admixture event occurred between populations after the split (scenarios S+SC+SB+SCB; 2,013 of 3,000 RF-trees; posterior probability = 0.700). When considering the twelve scenarios separately, the highest classification vote was for scenario SCB (1,521 of 3,000 RF-trees), which congruently excludes a genetic admixture event and includes a population size contraction in the ancestral population as well as a bottleneck event at the time of divergence in the *S. g. flaviventris* subspecies. The posterior probability of scenario SCB averaged 0.564 over the ten replicate analyses (Table 2). Table S1.1 (Supplementary Material S1) shows that only two other scenarios (SB and SCBA_U_) obtained at least 5% of the votes. The scenario SB included only a single bottleneck event in *S. g. flaviventris* (mean of 15.7% of votes) and the scenario SCBA_U_ included a bottleneck event in *S. g. flaviventris*, a population size contraction in the ancestral population and an unidirectional genetic admixture event from *S. g. gregaria* into *S. g. flaviventris* (mean of 10.8% of votes). All other scenarios obtained less than 5% of the votes and were hence even more weakly supported.

**Table 2.**
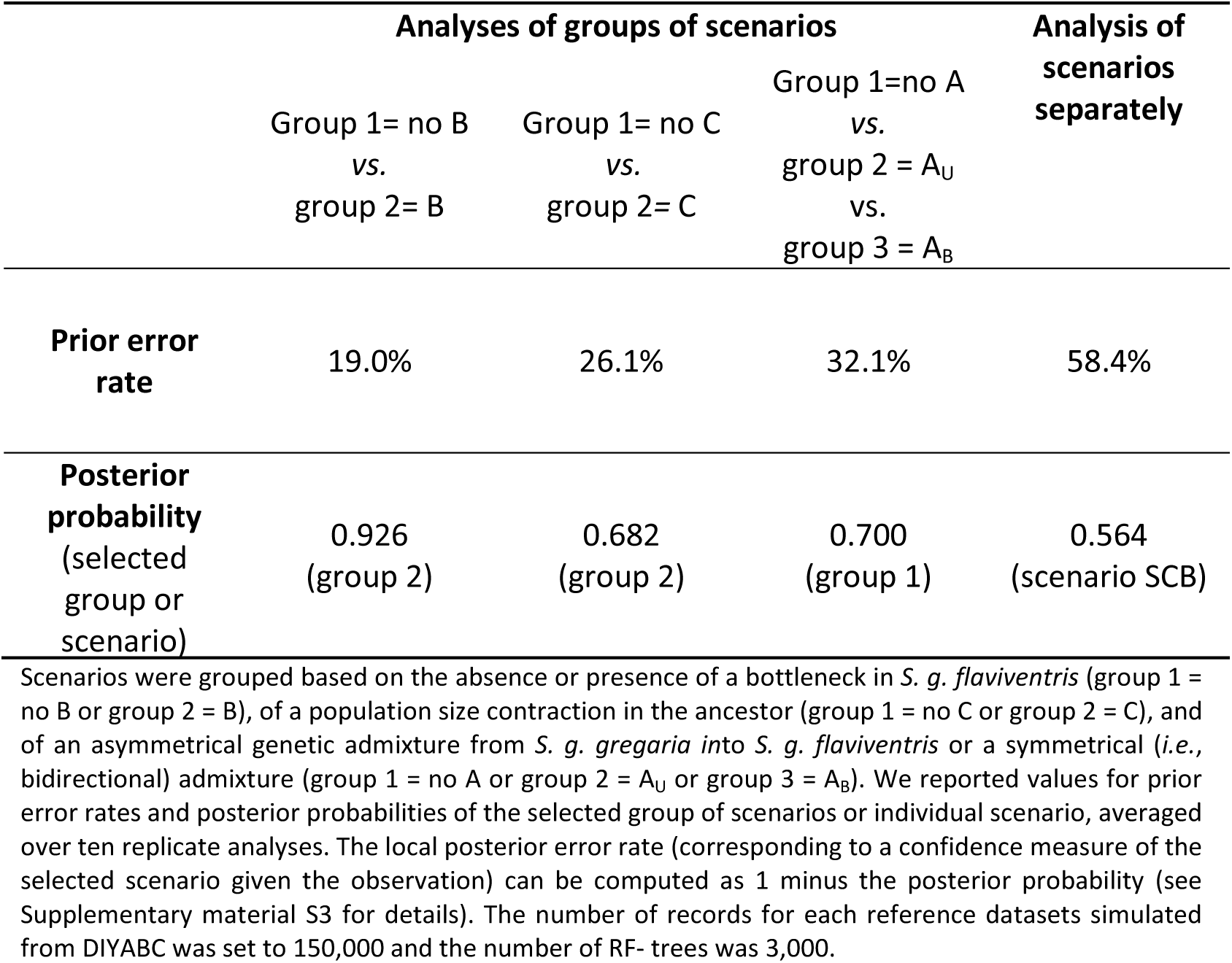
Scenario choice when analyzing groups of scenarios or scenarios separately.

|  | Analyses of groups of scenarios |  |  | Analysis of scenarios separately |
| --- | --- | --- | --- | --- |
|  | Group 1= no B<br>vs.<br>group 2= B | Group 1= no C<br>vs.<br>group 2= C | Group 1=no A<br>vs.<br>group 2 = A<sub>U</sub><br>vs.<br>group 3 = A<sub>B</sub> |  |
| **Prior error rate** | 19.0% | 26.1% | 32.1% | 58.4% |
| **Posterior probability**<br>(selected group or scenario) | 0.926<br>(group 2) | 0.682<br>(group 2) | 0.700<br>(group 1) | 0.564<br>(scenario SCB) |

Scenarios were grouped based on the absence or presence of a bottleneck in *S. g. flaviventris* (group 1 = no B or group 2 = B), of a population size contraction in the ancestor (group 1 = no C or group 2 = C), and of an asymmetrical genetic admixture from *S. g. gregaria* into *S. g. flaviventris* or a symmetrical (*i.e.*, bidirectional) admixture (group 1 = no A or group 2 = A<sub>U</sub> or group 3 = A<sub>B</sub>). We reported values for prior error rates and posterior probabilities of the selected group of scenarios or individual scenario, averaged over ten replicate analyses. The local posterior error rate (corresponding to a confidence measure of the selected scenario given the observation) can be computed as 1 minus the posterior probability (see Supplementary material S3 for details). The number of records for each reference datasets simulated from DIYABC was set to 150,000 and the number of RF- trees was 3,000.

We found that the posterior error rates (*i*.*e*., 1 minus the posterior probabilities) were lower than the prior error rates for the analyses considering either groups of scenarios based on the presence (or not) of a bottleneck in *S. g. flaviventris* (*i*.*e*., 7.4% *versus* 19.0%) or the scenarios separately (*i*.*e*., 43.6% *versus* 58.4%). For other groups of scenarios, the discrimination power was similar at both the global (prior error rates) and local (posterior error rates) scales, with values ranging from 26.1% to 32.1% (Table 2). Altogether, these results indicate that the observed dataset belongs to a region of the data space where the power to discriminate among scenarios is higher than the global power computed over the whole prior data space, and that the presence or absence of a bottleneck in *S. g. flaviventris* is the demographic event with the most robust prediction in our ABC-RF treatments. These results can be visually illustrated by the projection of the reference table datasets and the observed one on a single (when analyzing pairwise groups of scenarios) or on the first two linear discriminant analysis (LDA) axes (when analyzing the twelve scenarios considered separately) (Figure S1.1, Supplementary Material S1).

Figure S1.2, Supplementary Material S1, illustrates how RFs automatically rank the summary statistics according to their level of information. It shows that the number and set of most informative statistics is different depending on the comparisons (groups of scenarios or individual scenarios). Two sample statistics that measure the amount of genetic variation shared between populations (*F*_ST_, DM2 and LIK) were among the most informative when discriminating among groups of scenarios including or not an admixture event. For groups of scenarios differing by population size variation events, statistics summarizing variation between the two subspecies samples (*F*_ST_ and DM2 for the bottleneck event in *S. g. flaviventris*; DAS and LIK for the population size contraction in the ancestral population) and statistics summarizing genetic variation within subspecies samples (mean expected heterozygosity and mean number of alleles for both population size variation events) were among the most discriminative ones. Only fifteen single sample statistics were not informative (according to their position relatively to the noise statistics added to our treatments) when considering the twelve individual scenarios separately. Most of those non informative statistics were associated to the set of transcribed microsatellites (Figure S1.3, Supplementary Material S1).

### Parameter estimation under the best evolutionary scenario

Figure 3a shows point estimates with 90% credibility intervals of the posterior distribution of the divergence time between the two subspecies under the best supported scenario SCB. Our estimations point to a young age of subspecies divergence, with a median divergence time of 2.6 Ky and a 90% credibility interval of 0.9 to 6.6 Ky, when assuming an average of three generations per year (Roffey and Magor 2003; see Table 3 and Table S1.2, Supplementary Material S1, for details). The accuracy of divergence time estimation was almost similar at both the global and local scales (*i*.*e*., normalized mean absolute errors of 0.369 and 0.359, respectively; Table 3). Constraints on allele sizes in conjunction with high population-scaled mutation rates potentially strongly affect the linearity of the relationship between mutation accumulation and time of divergence estimated from microsatellite data. We thus evaluated the accuracy of ABC-RF estimation of the population divergence time as a function of the time scale, under scenario SCB. Analyses of simulated pseudo-observed datasets showed that the ABC-RF median estimate of divergence time reached a plateau for time scales ≥ 100,000 generations (Figure 4). Thus, the divergence time between *S. g. flaviventris* and *S. g. gregaria* estimated on our real microsatellite dataset (∼10,000 generations) is positioned within the period of linearity with time, well before reaching a plateau reflecting a saturation of genetic information at microsatellite markers. It is hence expected to represent a sensible estimation of the actual divergence time.

**Table 3.** Parameter estimation under the best supported scenario (scenario SCB)

|  | Posterior values |  |  | Prior values |  |  |
| --- | --- | --- | --- | --- | --- | --- |
|  | Median | 90% CI | NMAE | Median | 90% CI | NMAE |
| $t_s$ | 7,723 | 2,785<br>–<br>19,708 | 0.369 | 1,212 | 124<br>–<br>73,795 | 0.359 |
| $t_c / t_s$ | 2.75 | 1.11<br>–<br>35.47 | 0.596 | 12.17 | 1.24<br>–<br>762.26 | 1.077 |
| $N_f / N_g$ | 5.43 | 0.52<br>–<br>25.56 | 1.332 | 1.00 | 0.12<br>–<br>8.11 | 1.726 |
| $d_B / N_B$ | 1.06 | 0.49<br>–<br>2.41 | 0.323 | 1.00 | 0.13<br>–<br>7.57 | 0.299 |

$t_s$ : time of divergence between the two desert locust subspecies (in number of generations); $t_c$ : time of ancestral population size contraction; $N_f$ : stable effective population size of *S. g. flaviventris*; $N_g$ : stable effective population size of *S. g. gregaria*; $d_B$ : duration of the bottleneck event; $N_B$ : effective population size during the bottleneck event. For each evolutionary parameter, we reported posterior point estimates averaged over ten replicate analyses. CI: credibility interval. The number of records for each reference datasets simulated from DIYABC was set to 100,000 and the number of RF-trees was 2,000. Accuracy has been measured with the normalized mean absolute error (NMAE) which corresponds to the mean of the absolute difference between the point estimate of the parameter (here the median) and the (true) simulated value divided by the (true) simulated value. NMAE measures were computed using out-of-bag predictions either on the whole data space defined by the prior distributions (prior NMAE) or conditionally to the observed dataset (posterior NMAE); see Supplementary material S3 for details.

**Figure 3.**
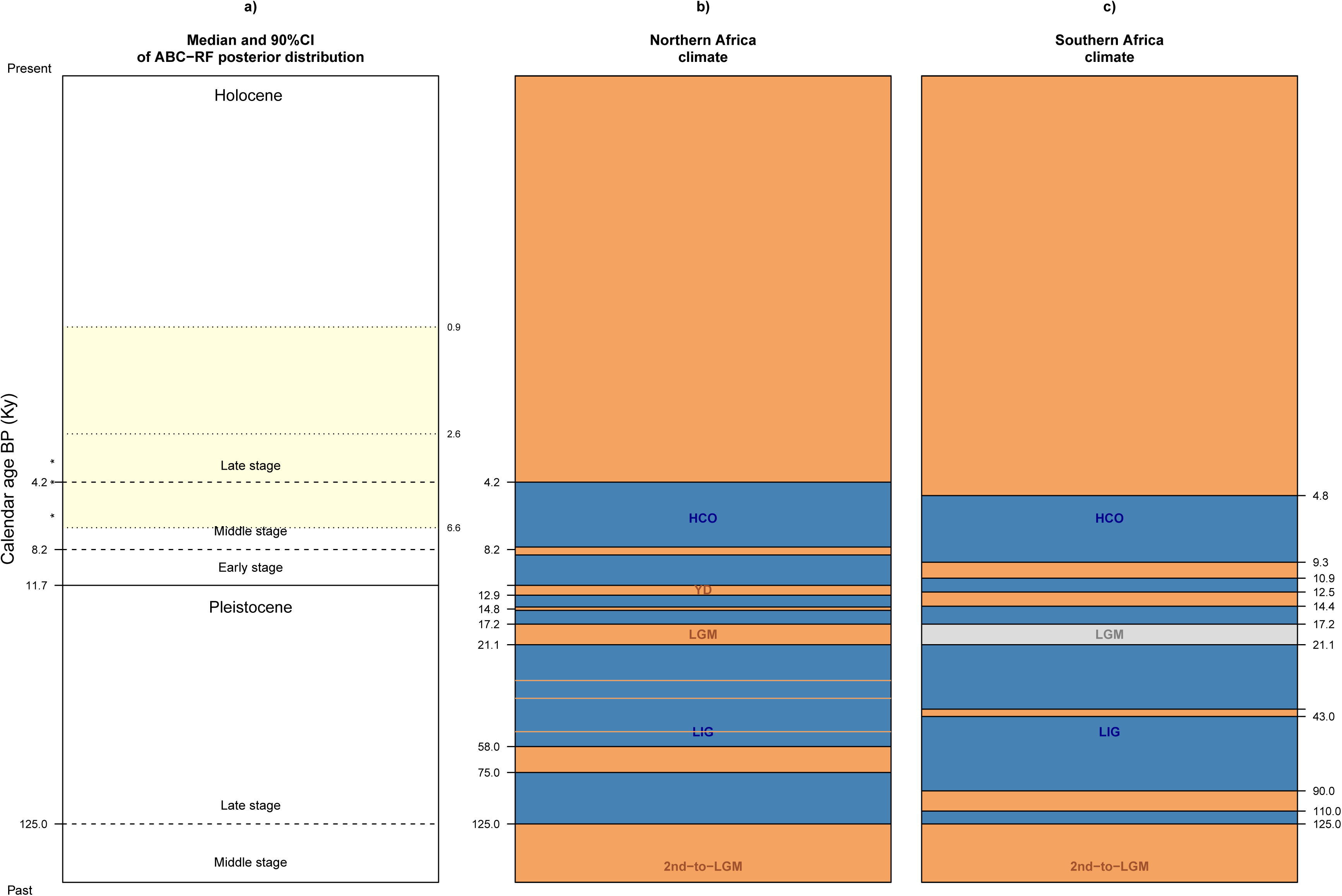
Divergence time between S. g. gregaria and S. g. flaviventris inferred under the best supported scenario (scenario SCB) a) in relation to bioclimatic changes in Northern b) and Southern Africa c). a) Dashed and solid lines represent the formal subdivision of the Holocene and Pleistocene epochs (Walker et al. 2012). Dotted lines with labels on the right side are the median value and 90% credibility interval of the posterior density distributions of the divergence time (assuming an average of three generations per year; Roffey and Magor 2003). Asterisks refer to earliest archeological records of the desert locust. In the Algerian Sahara, remains of locusts were found in a special oven dating back to about 6Ky ago, in the rock shelter of Tin Hanakaten (Aumassip 2002). In Egypt, locusts were depicted on daggers of the pharaoh Ahmose, founder of the Eighteenth Dynasty (about 3.5 Ky ago) (Malek 1997) and, at Saqqara, on tombs of the Sixth Dynasty (about 4.2 to 4.4 Ky ago) that is thought to have felt with the impact of severe droughts (Meinzingen 1993). b-c) Climatic episodes include major cycles and additional transitions of aridity (sandy brown) and humidity (steel blue). The grey coloration means that there is debate on the climatic status of the period (arid versus humid). HCO: Holocene Climatic Optimum; YD: Younger Dryas; LGM: Last Glacial Maximum; LIG: Last Inter Glacial. Delimitations of climatic periods were based on published paleoclimatic inferences from geological sediment sequences (e.g., eolian deposition, oxygen isotope data) and biological records (e.g., pollen or insect fossils assemblages) from marine cores or terrestrial lakes. References are Bond et al. (1997), Guo et al.(2000), Kröpelin et al. (2008), Roberts et al. (1993) and van Andel and Tzedakis (1996) for northern Africa, and Talma and Vogel (1992), Stokes et al. (1997), and Shi et al. (1998) for southern Africa. See also Gasse (2000) for a review.

**Figure 4.**
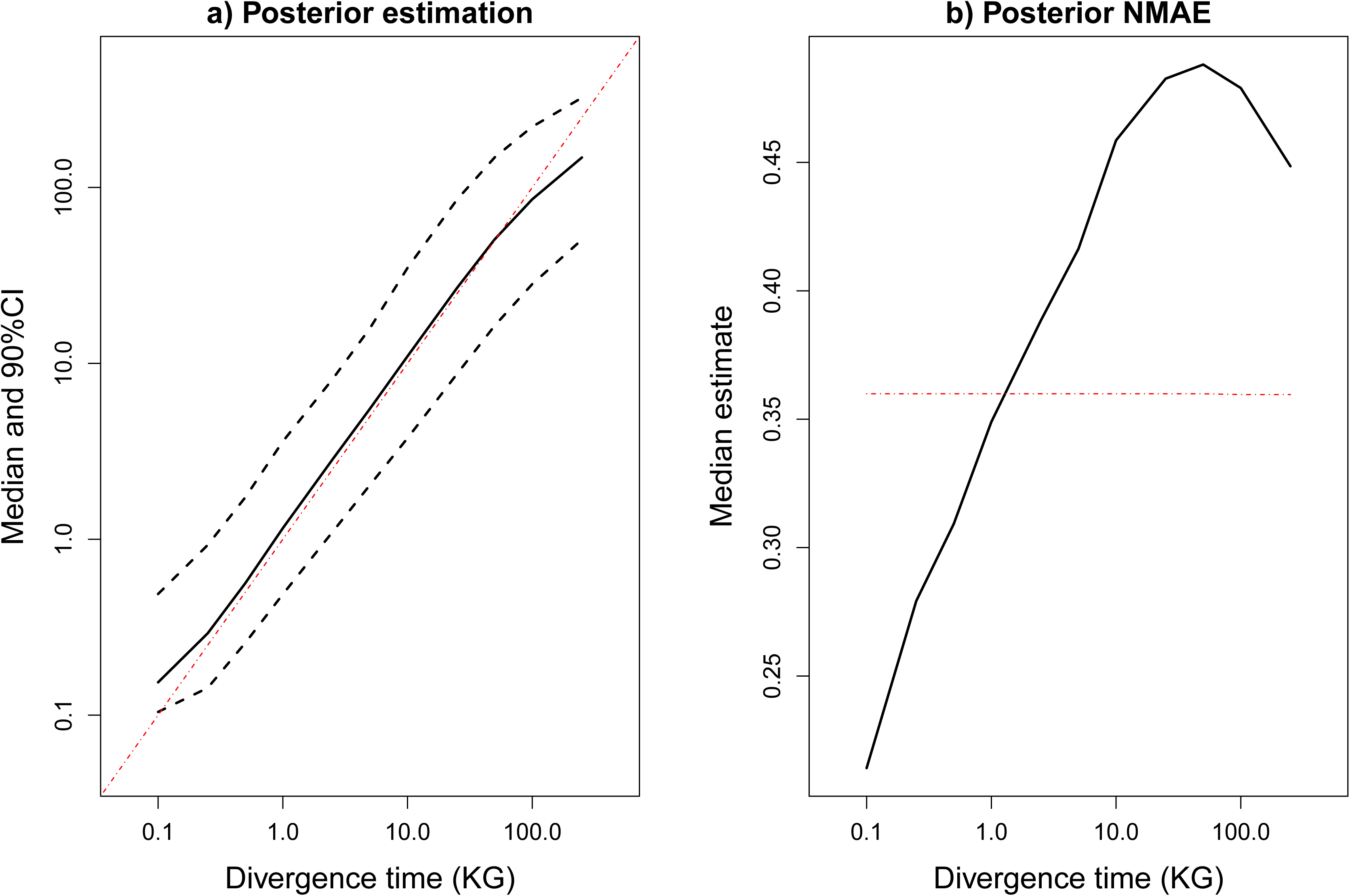
Median and 90% credibility interval a) and local accuracy b) of ABC-RF posterior distributions of the divergence time under the best supported scenario (scenario SCB) as a function of the time scale. Simulated pseudo-observed datasets (5,000 per divergence time) were generated for fixed divergence time values of 100; 250; 500; 1,000; 2,500; 5,000; 10,000; 25,000; 50,000; 100,000; and 250,000 generations (cf. x-axis with a log-scale). a) The estimated median (plain lines) and 90% credibility interval (90% CI; dashed lines), averaged over the 5,000 datasets, are represented (y-axis). b) The local accuracy is measured using out-of-bag predictions and the normalized mean absolute error (NMAE). See Supplementary Material S3 for details.

Using the median as a point estimate, we estimated that the population size contraction in the ancestor could have occurred at a time approximately three times older than the divergence time between the subspecies (cf. estimation of the parameter ratio *t*_*C*_ */ t*_*S*_ in Table 3). Estimations of the ratio of stable effective sizes of the *S. g. gregaria* and *S. g. flaviventris* populations (*i*.*e*., *N*_*f*_ / *N*_*g*_) showed large 90% credibility intervals and include the rate value of 1 (Table 3). Accuracy analysis indicates that our genetic data withhold little information on the composite parameter *N*_*f*_ / *N*_*g*_ (cf. the high associated NMAE values in Table 3). The bottleneck intensity during the colonization of south-western Africa (corresponding to the composite parameter *d*_*B*_ / *N*_*B*_) shows the highest accuracy of estimation among the parameters of interest (cf. the lowest associated NMAE values in Table 3). The median of 1 and the 90% credibility interval of 0.5 to 2.4 support a strong to moderate bottleneck event (Table 3).

For the time since divergence between the two subspecies, the most informative statistics corresponded to the expected heterozygosity computed within the *S. g. flaviventris* sample and the mean index of classification from *S*. *g. flaviventris* to *S. g. gregaria* (Figure S1.4, Supplementary Material S1). The addition of noise variables in our treatments showed that most statistics characterizing genetic variation within the *S. g. gregaria* sample were not informative. The most informative summary statistics were different depending on the parameter of interest (results not shown).

## Discussion

### A young age of subspecific divergence

With a 90% credibility interval of the posterior density distribution of the divergence time at 0.9 to 6.6 Ky, our ABC-RF analyses clearly point to a divergence of the two desert locust subspecies occurring during the present Holocene geological epoch (0 to 11.7 Ky ago; Figure 3a). The posterior median estimate (2.6 Ky) and interquartile range (1.8 to 3.7 Ky) postdated the middle-late Holocene boundary (4.2 Ky). The latter past time boundary corresponds to the last transition from humid to arid conditions in the African continent (Figure 3b). This increasing aridity was shown to be a progressive change, with a concomitant maximum in northern and southern Africa at around 4.0 to 4.2 Ky ago, where aridity caused a contraction of the forest at its northern and southern peripheries without affecting its core region (Guo *et al*. 2000; Maley *et al*. 2018). Interestingly, the earliest archeological records of the desert locust found in Tin Hanakaten (Algeria) and Saqqara (Egypt) archaeological sites date back to this period (Figure 3b; Meinzingen 1993; Malek 1997; Aumassip 2002). Pollen records also showed that during this period the plant community was dominated by the desert and semi-desert taxa found today, including some species of prime importance for the current ecology of the desert locust (Kröpelin *et al*. 2008, Shi *et al*. 1998, Duranton *et al*. 2012). Then, the past 4 Ky are thought to have been under environmental stability and as dry as at present. One can therefore reasonably assume that, at the inferred divergence time between the two locust subspecies, the connectivity between the two African hemispheres was still limited by the moist equator, in particular at the west, and by the savannahs and woodlands of the eastern coast (Figure 1c). Consequently, contrary to most phylogeographic studies on other African arid-adapted species (Atickem *et al*. 2018, Moodley *et al*. 2018), it is unlikely that the rather ancient Quaternary climatic history explained the Southern range extension of the desert locust.

Recent geological and palynological research has shown that a brief fragmentation of the African primary forest occurred during the Holocene interglacial from 2.5 Ky to 2.0 Ky ago (reviewed in Maley *et al*. 2018). This forest fragmentation period is characterized by relatively warm temperatures and a lengthening of the dry season rather than an arid climate. Although this period does not correspond to a phase of general expansion of savannas and grasslands, it led to the opening of the Sangha River Interval (SRI) in the core of the tropical forest in Central Africa (see Fig. 1 in Maley *et al*. 2018). The SRI corresponds to a 400 km wide (14–18° E) open strip composed of savannas and grasslands dividing the rainforest in a north-south direction. The SRI corridor is thought to have facilitated the southern migration of Bantu-speaking pastoralists, along with cultivation of the semi-arid sub-Saharan cereal, pearl millet, *Pennisetum glaucum* (Schwartz 1992; Bostoen *et al*. 2015). The Bantu expansion took place between approximately 5 and 1.5 Ky ago and reached the southern range of the desert locust, including northern Namibia for the Western Bantu branch and southern Botswana and eastern South Africa for the Eastern Bantu branch (Vansina 1995). We cannot exclude that the recent subspecific distribution of the desert locust has been mediated by this recent ecological disturbance, which included a north-south corridor of open vegetation habitats and the diffusion of agricultural landscapes through the Bantu expansion. The progressive reappearance of forest vegetation 2 Ky ago would have then led to the present-day isolation and subsequent genetic differentiation of the new southern populations from northern parental populations. The level of climatic and habitat favorability of the SRI environmental disturbance to the species, and thereby the likelihood of a south-eastern colonization through the SRI corridor, remains however to be evaluated in line with further data on this period.

### On the role of dispersal on subspecific divergence

Our ABC-RF results indicate that a demographic bottleneck (*i*.*e*., a strong transitory reduction of effective population size) occurred in the nascent southern subspecies of the desert locust. The high posterior probability value (92.6%) shows that this evolutionary event could be inferred with strong confidence. This result is compatible with the abovementioned colonization hypothesis if the proportion of suitable habitats for the desert locust in the SRI corridor was low, strongly limiting the carrying capacity during the time for range expansion. This scenario reduces, at the same time, the likelihood of a successful colonization through the SRI corridor. A more plausible alternative for a bottleneck event in *S. g. flaviventris* is a southern colonization of Africa through an exceptional long-distance migration event. In winged migratory species, movements are assisted by high velocity winds and may ascend to high altitudes (2000 m in the desert locust; Uvarov 1977) (Pedgley *et al*. 1995). Although largest insects exert some control over their direction of migration by flying actively, or at least gliding within (Pedgley *et al*. 1995), accidental displacements in wind directions markedly different from that of seasonal dominant winds that assist migrations are likely. Such accidental displacements at long-distance (*i*.*e*., a few to several thousand km) were recorded in a number of migratory species, such as cicadas, butterflies, moths and locusts (Pedgley *et al*. 1995; Lorenz, 2009).

To survive in its erratic arid and semi-arid habitat, the desert locust migrates downwind to reach areas where rain has recently fallen and exploit newly available resources. The dynamics of prevailing winds and pressure over Africa predicts the likelihood of a south-eastern transport of locusts (Nicholson 1996, Waloff and Pedgley 1986). In northern Africa, at least since 2.7Ky, the strong northeast trade winds bring desert locust swarms equatorward in the moist intertropical convergence zone (Figures 1a-b; Kröpelin *et al*. 1998). Most winds are westerlies (Figures 1a-b), nevertheless easterly winds flow parallel to the eastern coast of Africa in northern winter (*e*.*g*., January; Figure 1b). In southern Africa, winds blow mostly from the north-east toward the extant south-western distribution of the desert locust in southern winter (*e*.*g*., August; Figure 1a). In agreement with this, southward movements of desert locust have been documented along the eastern coast of Africa, in southern Tanzania during the plagues of 1926-1934, 1940-1948 and 1949-1963 (Waloff 1976), and even in Mozambique in January 1945 during the peak of the major plague of 1940-1948 (Waloff 1966).

Furthermore, most travels off the range listed in the history of the desert locust were associated with plague events (Waloff 1976), with other records including Portugal, the British Isles and the famous trans-Atlantic crossing observed in October 1988, where large numbers of locusts landed in the Caribbean islands (Richardson and Nemeth, 1991). Plagues typically culminate several years of above average rainfall, resulting in abundant vegetation that supports both cumulative locust population growth and full development of gregarious characteristics (Richardson and Nemeth, 1991). The gigantic numbers (over a billion) of swarming locusts may facilitate the success of long-distance migrations, in spite of high mortality. In addition, flight capacity and endurance of the gregarious phase are remarkable, with swarms of winged adults regularly travelling up to hundreds km in a day, in case of sudden lack of resources (Roffey and Magor 2003). Therefore, out-of-range displacements of the desert locust may be explained by the synergistic effects of exceptional environmental conditions (*e*.*g*., unusual winds in strength or direction, rain favorable to plague) and peculiar biology of the gregarious phase of this species (*e*.*g*., swarming behavior, huge numbers of dispersers).

In conclusion, dynamics of dominant winds in subtropical desert, historical records of exceptional migrations in the desert locust, and the very robust prediction for the presence of a bottleneck in *S. g. flaviventris* right after divergence by our ABC-RF treatments altogether support that a single or a few swarm(s) from the central region of the desert locust range sourced during a plague the colonization of south-western Africa, suggesting a role of dispersal in the disjoint distribution and divergence of desert locust populations. Interestingly, an exceptional trans-Atlantic flight of the desert locust from Africa to South America was revealed to give rise to the radiation of some 50 *Schistocerca* species in the western hemisphere (Lovejoy *et al*., 2006).

### On the influence of climatic cycles

It may appear surprising, at least at first sight, that the southern colonization of the desert locust did not occur during one of the major glacial episodes of the last Quaternary cycle, since these periods are characterized by a more continuous range of the desert locust (see paleo-vegetation maps in Figures 1e-f). In particular, during the last glacial maximum (LGM, −14.8 Ky to −26 Ky), the Sahara desert extended hundreds of km further South than at present and annual precipitation were lower (i.e. ∼200–1,000 mm/year). Several hypotheses explain why our evolutionary scenario choice procedure provided low support to the possibility of a birth of the locust subspecies *S. g. flaviventris* at older periods. First, we cannot exclude that our microsatellite genetic data allow making inferences about the last colonization event only. Error rates at both local and global scales for the choice of scenario groups including or not a genetic admixture event after the split indicated that our discrimination power to infer this specific evolutionary event was poor (*i*.*e*., local and global error rates of 32.1% and 30.0%, respectively). The recent north-to-south colonization event selected by our ABC-RF treatment may hence have blurred traces of older colonization events.

Second, while there is large evidence that much of Africa was drier during the last glacial phase, this remains debated for south-western Africa (see the gray coloration in Figure 3b). Some climate models show that at least some parts of this region, such as the Kalahari Desert, may have experienced higher rainfall than at present (Cockcroft *et al*. 1987; Ganopolski *et al*. 1998; Chase and Meadows 2007). Such regional responses to glacial cycles may have prolonged until the middle Holocene. In particular, the northern Younger Dryas (*i*.*e*., −12.9 to −11.7 Ky) can be correlated only partly with an arid period in the southern hemisphere (*i*.*e*., −14.4 to −12.5 Ky). Such older climate episodes in antiphase between hemispheres (see the sandy brown coloration in Figure 3b) may have prevented from either a successful north-to-south migration event or a successful establishment and spread in the new southern range.

Third, although semi-desert and desert biomes were more expanded than at present during the LGM, extreme aridity and lowered temperatures may have actually been unfavorable to the species. For instance, mean temperatures lowered by 5 to 6°C in both southern-western Africa (Stute and Talma 1997) and Central Sahara (Edmunds *et al*. 1999). The maintenance of desert locust populations depends on the proximity of areas with rainfalls at different seasons or with the capacity to capture and release water. For instance, in the African northern range, breeding success of locust populations relies on seasonal movements between the Sahel-Saharan zones of inter-tropical convergence, where the incidence of rain is high in summer, and the Mediterranean-Saharan transition zone, with a winter rainfall regime (Rainey and Waloff 1951). In addition, adult migration and nymphal growth of the desert locust are dependent upon high temperature (Roffey and Magor 2003). It is hence possible then that the conjunction of hyper-aridity with intense cold could not easily support populations of the desert locust, despite the high extent of their migrations. Interestingly, climatic reconstructions during the Last Glacial Maximum (LGM) showed dramatic decreases of mean temperature throughout all of Africa and of precipitation in the Sahelian region (see Figs. S2.3 and S2.4, Supplementary Material S2). Accordingly, species distribution modelling showed a LGM distribution for the desert locust very similar to its current distribution (Figs. S2.5, S2.6 and S2.7, Supplementary Material S2), excluding the hypothesis of a more continuous range of the species at this time.

While ABC-RF analyses did not support that the Quaternary climatic history explained the subspecific divergence in the desert locust, they provided evidence for the occurrence of a large contraction of the size of the ancestral population preceding the divergence. Using the median as a point estimate, we estimated that the population size contraction in the ancestor could have occurred at a time about three fold older than the divergence time between the subspecies. This corresponds to the African humid period in the early and middle stages of the Holocene, though the large credibility interval also included the last interglacial period of the Pleistocene (Figure 3b). Such population size contraction was likely induced by the severe(s) contraction(s) of deserts that prevailed prior the estimated divergence between the two subspecies. Interestingly, these humid periods were more intense and prolonged in northern Africa, which corresponded to the presumed center of origin of the most recent common desert locust ancestor (Scott 1993; Partridge 1997; Shi *et al*. 1998).

### Statistical advances by means of ABC Random Forest

To our knowledge, the present study is the first one using recently developed ABC-RF algorithms (Raynal *et al*. 2019) to carry out inferences about parameter estimation on a real multi-locus microsatellite dataset (for scenario choice see for example Rougemont *et al*. 2016 and Fraimout *et al*. 2017). When compared with various ABC solutions, this new RF method offers several advantages: a significant gain in terms of robustness to the choice of the summary statistics, independence from any type of tolerance level, and a good trade-off in terms of quality of point estimator precision of parameters and credible interval estimations for a given computing time (Raynal *et al*. 2019).

The RF approach that we used here includes three novelties in statistical analyses that were particularly useful for reconstructing the evolutionary history of the divergence between *S. g. gregaria* and *S. g. flaviventris* subspecies. First, our ABC-RF statistical treatments benefited from the incorporation of previous estimations of mutation rates and allele size constraints for the microsatellite loci used in this study (for details see the Materials and methods section *Microsatellite dataset, mutation rate and mutational model*). Microsatellite mutation rate and pattern of most eukaryotes remains to a large extent unknown, and, to our knowledge, the present study is a rare one where independent information on mutational features was incorporated into the microsatellite prior distributions. Second, given the low RF computing time, we could simulate large pseudo-observed datasets to compute error measures conditionally to a subset of fixed divergence time values chosen to cover the entire prior interval. In this way, we showed that posterior distributions for estimation of divergence time between the two subspecies accurately reflected true divergence time values. Third, because error levels may differ depending on the location of an observed dataset in the prior data space (*e*.*g*., Pudlo *et al*. 2016), prior-based indicators are poorly relevant, aside from their use to select the best classification method and set of predictors, here our summary statistics. Therefore, in addition to global prior errors, we computed local posterior errors, conditionally to the observed dataset. The latter errors measure prediction quality exactly at the position of the observed dataset. For parameter estimation accuracy, we propose an innovative way to approximate local posterior errors, relying partly on out-of-bag predictions and implemented in a new version of the R library abcrf (version 1.8) available on R CRAN (see the Supplementary Material S3 for a detailed description). Such statistical advances are of general interest and will be useful for any statistical treatments of massive simulation data, including for inferences using single nucleotide polymorphisms (*i*.*e*., SNPs) obtained from new generation sequencing technologies.

## Materials & Methods

### Formalization of evolutionary scenarios

To help formalize the evolutionary scenarios to be compared, we relied on maps of vegetation cover in Africa from the Quaternary Environment Network Atlas (Adams and Faure 1997). We considered the periods representative of arid maximums (YD and LGM; Figs. 1e-f), humid maximums (HCO; Fig. 1d), and present-day arid conditions (Fig. 1c), for which reliable vegetation reconstructions have been published. Desert and xeric shrubland cover fits well with the present-day species range during remission periods. Tropical and Mediterranean grasslands were added separately since the species inhabits such environments during outbreak periods only. The congruence between present maps of species distribution (Fig. 1a) and of open vegetation habitats (Fig. 1c) suggests that vegetation maps for more ancient periods could be considered as good approximations of the potential range of the desert locust in the past (but see section *On the influence of climatic cycles* in Discussion). Maps of vegetation cover during ice ages (Figs. 1e-f) show an expansion of open vegetation habitats (*i*.*e*., grasslands in the tropics and deserts in both the North and South of Africa) sufficient to make the potential range of the species continuous from the Horn of Africa in North-West to the Cape of Good Hope in the South. It is worth stressing that we also explored species distribution modelling for the HCO and LGM periods as an alternative to using vegetation as a surrogate for the locust range (detailed in Supplementary Material S2). However, distribution modelling provided a narrower set of alternative hypotheses than the vegetation-based scenarios mentioned above and are therefore not discussed any further.

Based on the above paleo-vegetation map reconstructions, we considered a set of alternative biogeographic hypotheses formulated into different types of evolutionary scenarios. First, we considered scenarios involving a more or less continuous colonization of southern Africa by the ancestral population from a northern origin. In this type of scenario, effective population sizes were allowed to change after the divergence event, without requiring any bottleneck event (*i*.*e*., without any abrupt and strong reduction of population size) right after divergence. Second, we considered the situation where the colonization of Southern Africa occurred through a single (or a few) long-distance migration event(s) of a small fraction of the ancestral population. This situation was formalized through scenarios that differed from a continuous colonization scenario by the occurrence right after divergence of a bottleneck event in the newly founded population (*i*.*e*., *S. g. flaviventris*), which was modelled through a limited number of founders during a short period.

Because the last Quaternary cycle includes several arid climatic periods, including the intense punctuation of the Younger Dryas (YD) and the last glacial maximum (LGM), we also considered scenarios that incorporated the possibility of a discrete genetic admixture event, either bidirectional or unidirectional from *S. g. gregaria* into *S. g. flaviventris*. Since previous tests based on simulated data showed a poor power to discriminate between a single *versus* several admixture events (results not shown), we considered only scenarios including a single admixture event.

Finally, at interglacial humid maximums, the map of vegetation cover showed a severe contraction of deserts, which were nearly completely vegetated with annual grasses and shrubs and supported numerous perennial lakes (Fig. 1d; deMenocal *et al*. 2000). We thus considered the possibility that vegetation-induced contractions of population sizes have pre-dated the separation of the two subspecies. Hence, whereas so far scenarios involved a constant effective population size in the ancestral population, we formalized alternative scenarios in which we assumed that a long population size contraction event occurred into the ancestral population.

Combining the presence or absence of the three above-mentioned key evolutionary events (a bottleneck in *S. g. flaviventris*, a bidirectional or unidirectional genetic admixture from *S. g. gregaria* into *S. g. flaviventris*, and a population size contraction in the ancestral population) allowed defining a total of twelve scenarios, that we compared using ABC-RF. The twelve scenarios with their historical and demographic parameters are graphically depicted in Figure 2. All scenarios assumed a northern origin for the common ancestor of the two subspecies and a subsequent southern colonization of Africa. This assumption is supported by recent mitochondrial DNA data showing that *S. g. gregaria* have higher levels of genetic diversity and diagnostic bases shared with outgroup and congeneric species, whereas *S. g. flaviventris* clade was placed at the apical tip within the species tree (Chapuis *et al*. 2016). In agreement with this assumption, preliminary analyses based on observed data showed a low support for a southern origin for the common ancestor of the two subspecies and a subsequent northern colonization of Africa (results not shown).

All scenarios considered three populations of stable effective population sizes *N*_*f*_ for *S. g. flaviventris, N*_*g*_ for *S. g. gregaria*, and *N*_*a*_ for the ancestral population, with *S*. *g. flaviventris* and *S. g. gregaria* diverging *t*_*S*_ generations ago from the ancestral population. The bottleneck event which potentially occurred into *S. g. flaviventris* was modelled through a limited number of founders *N*_*B*_ during a short period *d*_*B*_. The potential unidirectional genetic admixture into *S. g. flaviventris* occurred at a time *t*_*A*_, with a proportion *r*_*AU*_ of genes of *S. g. gregaria* origin. In the case of a bidirectional genetic admixture, still occurring at a time *t*_*A*_, the proportion of *S. g. gregaria* genes entering into the *S. g. flaviventris* population was *r*_*AB*_ and the proportion of *S. g. flaviventris* genes entering into the *S. g. gregaria* population was 1-*r*_*AB*_. The potential population size contraction event occurred into the ancestral population at a time *t*_*C*_, with an effective population size *N*_*C*_ during a duration *d*_*C*_.

### Prior setting for historical and demographical parameters

Prior values for time periods between sampling and admixture, divergence and/or ancestral population size contraction events (*t*_C_, *t*_S_ and *t*_A_, respectively) were drawn from log-uniform distributions bounded between 100 and 500,000 generations, with *t*_C_ > *t*_S_ > *t*_A_. Assuming an average of three generations per year (Roffey and Magor 2003), this prior setting corresponds to a time period that goes back to the second-to-latest glacial maximum (150 Ky ago) (de Vivo and Carmignotto 2004, deMenocal *et al*. 2000). Preliminary analyses showed that assuming a uniform prior shape for all time periods (instead of log-uniform distributions) do not change scenario choice results, with posterior probabilities only moderately affected, and this despite a substantial increase of out-of-bag prior error rates (*e*.*g*., + 50% when considering the twelve scenarios separately; Table S4.1, Supplementary Material S4). Analyses of simulated pseudo-observed datasets (pods) showed that assuming a uniform prior rather than a log-uniform prior for time period parameters would have also biased positively the median estimate of the divergence time and substantially increased its 90% credibility interval (Figure S4.1 and Table S4.2, Supplementary Material S4). Using a log-uniform distribution remains a sensible choice for parameters with ranges of values covering several if not many log-intervals, as doing so allows assigning equal probabilities to each of the log-intervals.

We used uniform prior distributions bounded between 1−10^4^ and 1−10^6^ diploid individuals for the different stable effective population sizes *N*_*f*_, *N*_*g*_ *and N*_*a*_ (Chapuis *et al*. 2014). The admixture rate (*i*.*e*., the proportion of *S. g. gregaria* genes entering into the *S. g. flaviventris* population), was drawn from a uniform prior distribution bounded between 0.05 and 0.50 for a unidirectional event (*r*_*AU*_) and between 0.05 and 0.95 for a bidirectional event (*r*_*AB*_). We used uniform prior distributions bounded between 2 and 100 for both the numbers of founders (in diploid individuals) and durations of bottleneck events (in number of generations). For the contraction event, we used uniform prior distributions bounded between 100 and 10,000 for both the population size *N*_*C*_ (in diploid individuals) and duration *d*_*C*_ (in number of generations). Assuming an average of three generations per year (Roffey and Magor 2003), such prior choice allowed a reduction in population size for a short to a relatively long period, similar for instance to the whole duration of the HCO (from 9 to 5.5 Ky ago) which was characterized by a severe contraction of deserts.

### Microsatellite dataset, mutation rate and mutational model

We carried out our statistical inference on the microsatellite dataset previously published in Chapuis *et al*. (2016). The 23 microsatellite loci genotyped in that dataset were derived from either genomic DNA (14 loci) or messenger RNA (9 loci) resources, and were hereafter referred to as untranscribed and transcribed microsatellite markers (following Blondin *et al*. 2013). These microsatellites were shown to be genetically independent, free of null alleles and at selective neutrality (Chapuis *et al*. 2016). Previous levels of *F*_ST_ (Weir 1996) and Bayesian clustering analyses (Pritchard *et al*. 2000) among populations showed a weak genetic structuring within each subspecies (Chapuis *et al*. 2014, 2017). For each subspecies, we selected and pooled three population samples in order to ensure both a large sample size (*i*.*e*., 80 and 90 individuals for *S. g. gregaria* and *S. g. flaviventris*, respectively), while ensuring a non-significant genetic structure within each subspecies pooled sample, as indicated by non-significant Fisher’s exact tests of genotypic differentiation among the three initial population samples within subspecies and exact tests of Hardy-Weinberg equilibrium for each subspecies pooled sample (*i*.*e*., *P*-value > 0.05 when using Genepop 4.0; Rousset 2008). More precisely, the *S. g. gregaria* sample consisted in pooling the population samples 8, 15 and 22 of Chapuis *et al*. (2014) and the *S. g. flaviventris* sample included the population samples 1, 2 and 6 of Chapuis *et al*. (2017).

Mutations occurring in the repeat region of each microsatellite locus were assumed to follow a symmetric generalized stepwise mutation model (GSM; Zhivotovsky *et al*. 1997; Estoup *et al*. 2002). Prior values for any mutation model settings were drawn independently for untranscribed and transcribed microsatellites in specific distributions. Because allele size constraints exist at microsatellite markers, we informed for each microsatellite locus their lower and upper allele size bounds using values estimated in Chapuis *et al*. (2015), following the approach of Pollock *et al*. (1998) and microsatellite data from several species closely related to *S. gregaria* (Blondin *et al*. (2013). Prior values for the mean mutation rates 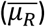 were set to the empirical estimates inferred from observation of germline mutations in Chapuis *et al*. (2015), *i*.*e*., 2.8−10^−4^ and 9.1−10^−5^ for untranscribed and transcribed microsatellites, respectively. The mutation rates for individual microsatellites were then drawn from a Gamma distribution with 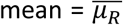 and shape = 0.7 (Estoup *et al*. 2001) for both types of microsatellites. We ensured that the chosen value of shape parameter generated the same inter-loci variance as estimated in Sun *et al*. (2012) from direct observations of thousands of human microsatellites. Prior values for the mean parameters of the geometric distributions of the length in number of repeats of mutation events 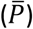 were set to the proportions of multistep germline mutations observed in Chapuis *et al*. (2015), *i*.*e*., 0.14 and 0.67 for untranscribed and transcribed microsatellites, respectively. The *P* parameters for individual loci were then standardly drawn from a Gamma distribution (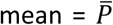 and shape = 2). We also considered mutations that insert or delete a single nucleotide to the microsatellite sequence. To model this mutational feature, we used the DIYABC default setting values (*i*.*e*., a uniform distribution bounded between [10^−8^, 10^−5^] for the mean parameter 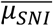 and a Gamma distribution (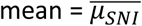 and shape = 2) for individual loci parameters; Cornuet *et al*. 2010; see also DIYABC user manual p. 13, http://www1.montpellier.inra.fr/CBGP/diyabc/).

### Analyses using ABC Random Forest

We used the software DIYABC v.2.1.0 (Cornuet *et al*. 2014) to simulate datasets constituting the so-called reference tables (*i*.*e*., records of a given number of datasets simulated using the scenario ID and the evolutionary parameter values sampled from prior distributions and summarized with a pool of statistics). Random-forest computations were then performed using a new version of the R library abcrf (version 1.8) available on the CRAN. This version includes all ABC-RF algorithms detailed in Pudlo *et al*. (2016), Raynal *et al*. (2019) and Estoup *et al*. (2018a) for scenario choice and parameter estimation, as well as several statistical novelties allowing to compute error rates in scenario choice and accuracy measures for parameter estimation (see details below). An overview of the ABC-RF methods used in the present paper is provided in Supplementary Material S3. Readers can also consult Pudlo *et al*. (2016), Rougemont *et al*. (2016), Fraimout *et al*. (2017), Estoup *et al*. (2018a,b) and Marin *et al*. (2018) for scenario choice, and Raynal *et al*. (2019) for parameter estimation to access to further detailed statistical descriptions, testing and applications of ABC-RF algorithms.

For scenario choice, the outcome of the first step of the ABC-RF statistical treatment applied to a given target dataset is a classification vote for each scenario which represents the number of times a scenario is selected in a forest of *n* trees. The scenario with the highest classification vote corresponds to the scenario best suited to the target dataset among the set of compared scenarios. This step also provides an error rate relevant to the entire prior sampling space, the global prior error. See the section *Global prior errors* in Supplementary Material S3 for details. The second RF analytical step provides a reliable estimation of the posterior probability of the best supported scenario. One minus such posterior probability yields the local posterior error associated to the observed dataset (see the section *Local posterior errors* in Supplementary Material S3). In practice, ABC-RF analyses were processed by drawing parameter values into the prior distributions described in the two previous sections and by summarizing microsatellite data using a set of 32 statistics (see Table 1 for details about such summary statistics) and the one LDA axis or eleven LDA axes (i.e. number of scenarios minus 1; Pudlo *et al*. 2016) computed when considering pairwise groups of scenarios or individual scenarios, respectively. We processed ABC-RF treatments on reference tables including 150,000 simulated datasets (*i*.*e*., 12,500 per scenario). Following Pudlo et al. (2016), we checked that 150,000 datasets was sufficient by evaluating the stability of prior error rates and posterior probabilities estimations of the best scenario on 50,000, 80,000 and 100,000 and 130,000 simulated datasets (Table S1.3, Supplementary Material S1). The number of trees in the constructed random forests was fixed to *n* = 3,000, as this number turned out to be large enough to ensure a stable estimation of the prior error rate (Figure S1.5, Supplementary Material S1). We predicted the best scenario and estimated its posterior probability and prior error rate over ten replicate analyses based on ten different reference tables.

In order to assess the power of our approach to infer each specific evolutionary event of interest, we first processed ABC-RF analyses by grouping scenarios based on the presence or absence of each type of evolutionary event that we identified as having potentially played a role in setting up the disjoint distribution of the two locust subspecies (*e*.*g*., Roux *et al*. 2016, Leroy *et al*. 2017 and Estoup *et al*. 2018a). We conducted ABC-RF treatments on three pairwise groups of scenario: groups of scenarios with *vs*. without a bottleneck in *S. g. flaviventris*, groups with *vs*. without a population size contraction in the ancestral population, and groups with *vs*. without an asymmetrical genetic admixture event from *S. g. gregaria* into *S. g. flaviventris*. We then conducted ABC-RF treatments on the twelve scenarios considered separately.

For parameter estimation, we conducted ten independent replicate RF treatments based on ten different reference tables for each parameter of interest (Raynal *et al*. 2019): the time since divergence, the ratio of the time of the contraction event into the ancestral population on the time since divergence, the intensity of the bottleneck event in the sampled *S. g. flaviventris* population (defined as the ratio of the bottleneck event of duration *d*_*B*_ on the effective population size *N*_*B*_) and the ratio of the stable effective population size of the two sampled populations. For each RF treatment, we simulated a total of 100,000 datasets for the selected scenario (drawing parameter values into the prior distributions described in the two previous sections and using the same 32 summary statistics). Following Raynal *et al*. (2019), we checked that 100,000 datasets was sufficient by evaluating the stability of the measure of accuracy on divergence time estimation using 50,000, 80,000 and 90,000 simulated datasets (Table S1.4, Supplementary Material S1). The number of trees in the constructed random forests was fixed to *n* = 2,000, as such number turned out to be large enough to ensure a stable estimation of the measure of divergence time estimation accuracy (Figure S1.6, Supplementary Material S1). For each RF treatment, we estimated the median value and the 5% and 95% quantiles of the posterior distributions. It is worth noting that we considered median values as the later provided more accurate estimations (according to out-of-bag predictions) than when considering mean values (results not shown). Accuracy of parameter estimation was measured using out-of-bag predictions and the normalized mean absolute error (NMAE). NMAE corresponds to the mean of the absolute difference between the point estimate (here the median) and the (true) simulated value divided by the simulated value (formula detailed in Supplementary Material S3).

Finally, because microsatellite markers tend to underestimate divergence time for large time scales due to allele size constraints, we evaluated how the accuracy of ABC-RF estimation of the time of divergence between the two subspecies was sensitive to the time scale. To this aim, we used DIYABC to produce simulated pseudo-observed datasets assuming fixed divergence time values chosen to cover the prior interval (100; 250; 500; 1,000; 2,500; 5,000; 10,000; 25,000; 50,000; 100,000; 250,000 generations) and using the best scenario. We simulated 5,000 of such test datasets for each of the eleven divergence time values. Each of these test dataset was treated using ABC-RF in the same way as the above target observed dataset.

## Data accessibility

Microsatellite genotypic data and R scripts for ABC-RF analyses are publicly available on the CIRAD Dataverse repository (https://dataverse.cirad.fr/) at https://doi.org/10.18167/DVN1/ZNSKGK.

## Supplementary material

Additional supporting information may be found in the online version of this article.

## Acknowledgements

This work was supported by research funds from the French Agricultural Research Centre for International Development (CIRAD), the project ANR-16-CE02-0015-01 (SWING), the INRA scientific department SPE (AAP-SPE 2016), and the Labex NUMEV (NUMEV, ANR10-LABX-20). The data used in this work were partly produced through the technical facilities of the Centre Méditerranéen Environnement Biodiversité, Montpellier. We are grateful to Pierre-Emmanuel Gay for assistance with maps, and Antoine Foucart, Gauthier Dobigny and Jean-Yves Rasplus for fruitful discussions. We are extremely grateful to the *Peer Community In Evolutionary Biology* recommenders (Concetta Burgarella and Takeshi Kawakami) and reviewers (Michael D Greenfield and two anonymous ones) for their wise and helpful comments on two earlier versions of the manuscript. The Peer Community In Evolutionary Biology recommendation of the version 4 of this preprint (along with peer-reviews and authors’ replies) is available at https://doi.org/10.24072/pci.evolbiol.100091.

## Conflict of interest disclosure

The authors of this preprint declare that they have no financial conflict of interest with the content of this article. Arnaud Estoup is one of the *PCI Evolutionary Biology* recommenders.

## Supplementary material online for the manuscript “A young age of subspecific divergence in the desert locust *Schistocerca gregaria* inferred by ABC Random Forest”

### Supplementary Material S1. Details on results from ABC-RF treatments

**Table S1.1.** Scenario choice for each of the ten replicate analyses.

| Reference table | Best scenario | Votes S | Votes SC | Votes SB | Votes SCB | Votes SA<sub>U</sub> | Votes SCA<sub>U</sub> | Votes SBA<sub>U</sub> | Votes SCBA<sub>U</sub> | Votes SA<sub>B</sub> | Votes SCA<sub>B</sub> | Votes SBA<sub>B</sub> | Votes SCBA<sub>B</sub> | Prior error rate | Posterior probability (best scenario) |
| --- | --- | --- | --- | --- | --- | --- | --- | --- | --- | --- | --- | --- | --- | --- | --- |
| 1 | scenario SCB | 0.004 | 0.033 | 0.171 | 0.513 | 0.011 | 0.022 | 0.040 | 0.094 | 0.017 | 0.041 | 0.024 | 0.031 | 0.583 | 0.572 |
| 2 | scenario SCB | 0.003 | 0.026 | 0.178 | 0.515 | 0.006 | 0.018 | 0.023 | 0.093 | 0.023 | 0.022 | 0.051 | 0.042 | 0.583 | 0.549 |
| 3 | scenario SCB | 0.003 | 0.024 | 0.196 | 0.497 | 0.010 | 0.019 | 0.030 | 0.101 | 0.025 | 0.033 | 0.036 | 0.024 | 0.583 | 0.545 |
| 4 | scenario SCB | 0.004 | 0.026 | 0.144 | 0.483 | 0.006 | 0.018 | 0.043 | 0.126 | 0.041 | 0.032 | 0.050 | 0.026 | 0.586 | 0.570 |
| 5 | scenario SCB | 0.004 | 0.025 | 0.126 | 0.537 | 0.007 | 0.017 | 0.030 | 0.117 | 0.017 | 0.021 | 0.063 | 0.036 | 0.582 | 0.607 |
| 6 | scenario SCB | 0.004 | 0.039 | 0.161 | 0.501 | 0.004 | 0.023 | 0.036 | 0.128 | 0.020 | 0.026 | 0.037 | 0.021 | 0.585 | 0.568 |
| 7 | scenario SCB | 0.014 | 0.021 | 0.136 | 0.536 | 0.005 | 0.022 | 0.029 | 0.099 | 0.022 | 0.044 | 0.037 | 0.035 | 0.585 | 0.588 |
| 8 | scenario SCB | 0.005 | 0.016 | 0.149 | 0.491 | 0.005 | 0.019 | 0.026 | 0.124 | 0.042 | 0.030 | 0.034 | 0.058 | 0.586 | 0.565 |
| 9 | scenario SCB | 0.003 | 0.036 | 0.151 | 0.506 | 0.009 | 0.019 | 0.017 | 0.113 | 0.049 | 0.024 | 0.034 | 0.040 | 0.584 | 0.551 |
| 10 | scenario SCB | 0.006 | 0.031 | 0.178 | 0.491 | 0.002 | 0.029 | 0.028 | 0.082 | 0.041 | 0.032 | 0.033 | 0.047 | 0.585 | 0.528 |
| Mean | scenario SCB | 0.005 | 0.028 | 0.159 | 0.507 | 0.007 | 0.021 | 0.030 | 0.108 | 0.030 | 0.030 | 0.040 | 0.036 | 0.584 | 0.564 |

**Table S1.2.**
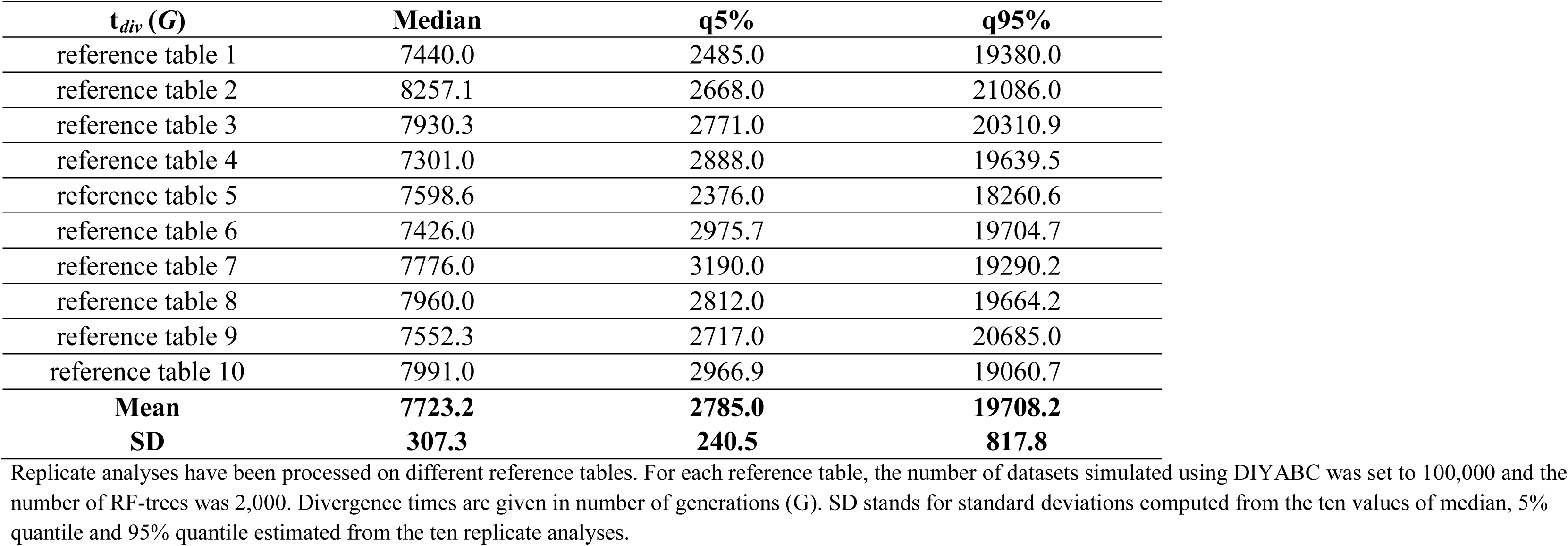
Estimation of the divergence time between *S. g. gregaria* and *S. g. flaviventris* for the ten replicate analyses processed under the best supported scenario (scenario SCB).

**Table S1.3.** Effect of the number of simulated datasets in the reference table on scenario choice.

| $n_{ref}$ | 50,000 | | 80,000 | | 100,000 | | 130,000 | | 150,000 | |
| --- | --- | --- | --- | --- | --- | --- | --- | --- | --- | --- |
|  | Mean | SD | Mean | SD | Mean | SD | Mean | SD | Mean | SD |
| **Votes S** | 0.006 | 0.002 | 0.005 | 0.002 | 0.006 | 0.003 | 0.006 | 0.004 | 0.005 | 0.003 |
| **Votes SC** | 0.030 | 0.015 | 0.031 | 0.009 | 0.029 | 0.006 | 0.028 | 0.006 | 0.028 | 0.007 |
| **Votes SB** | 0.155 | 0.025 | 0.155 | 0.028 | 0.165 | 0.033 | 0.164 | 0.036 | 0.159 | 0.022 |
| **Votes SCB** | 0.496 | 0.035 | 0.506 | 0.036 | 0.500 | 0.034 | 0.502 | 0.025 | 0.507 | 0.019 |
| **Votes SA<sub>U</sub>** | 0.006 | 0.003 | 0.007 | 0.004 | 0.006 | 0.004 | 0.006 | 0.004 | 0.007 | 0.003 |
| **Votes SCA<sub>U</sub>** | 0.025 | 0.005 | 0.023 | 0.004 | 0.020 | 0.005 | 0.022 | 0.006 | 0.021 | 0.008 |
| **Votes SBA<sub>U</sub>** | 0.033 | 0.009 | 0.031 | 0.007 | 0.033 | 0.008 | 0.030 | 0.009 | 0.030 | 0.016 |
| **Votes SCBA<sub>U</sub>** | 0.117 | 0.027 | 0.113 | 0.019 | 0.108 | 0.017 | 0.109 | 0.017 | 0.108 | 0.012 |
| **Votes SA<sub>B</sub>** | 0.031 | 0.009 | 0.030 | 0.011 | 0.027 | 0.010 | 0.029 | 0.010 | 0.030 | 0.008 |
| **Votes SCA<sub>B</sub>** | 0.032 | 0.012 | 0.029 | 0.007 | 0.029 | 0.009 | 0.031 | 0.012 | 0.030 | 0.011 |
| **Votes SBA<sub>B</sub>** | 0.039 | 0.014 | 0.039 | 0.013 | 0.043 | 0.014 | 0.042 | 0.013 | 0.040 | 0.011 |
| **Votes SCBA<sub>B</sub>** | 0.030 | 0.013 | 0.031 | 0.009 | 0.032 | 0.009 | 0.032 | 0.009 | 0.036 | 0.011 |
| **Prior error rate** | **0.594** | **0.002** | **0.589** | **0.002** | **0.587** | **0.001** | **0.585** | **0.001** | **0.584** | **0.001** |
| **Posterior probability**<br>**(best model)** | **0.567** | **0.037** | **0.562** | **0.034** | **0.555** | **0.034** | **0.563** | **0.028** | **0.564** | **0.023** |

**Table S1.4.** Effect of the number of simulated datasets in the reference table on posterior point estimation values a) and estimation accuracy b) of the divergence time between *S. g. gregaria* and *S. g. f laviventris* under the best supported scenario (scenario SCB).

| $n_{\text{ref}}$ | | 50,000 | | 80,000 | | 90,000 | | 100,000 | |
| --- | --- | --- | --- | --- | --- | --- | --- | --- | --- |
|  |  | Mean | SD | Mean | SD | Mean | SD | Mean | SD |
| **Posterior estimations** | Median | 7731.8 | 448.9 | 7691.8 | 357.5 | 7724.8 | 318.0 | 7723.2 | 307.3 |
|  | q5% | 2697.0 | 129.7 | 2706.8 | 235.2 | 2764.4 | 172.9 | 2785.0 | 240.5 |
|  | q95% | 20295.5 | 1763.6 | 19711.6 | 1451.3 | 19508.7 | 1264.1 | 19708.2 | 817.8 |

**b)**
| $n_{\text{ref}}$ | | 50,000 | 80,000 | 90,000 | 100,000 |
| --- | --- | --- | --- | --- | --- |
| **Accuracy measures** | Prior NMAE | 0.378 | 0.365 | 0.362 | 0.359 |
|  | Posterior NMAE | 0.375 | 0.370 | 0.365 | 0.369 |

The number of records in the reference datasets ( $n_{\text{ref}}$ ) simulated from DIYABC varied from 50,000 to 100,000. The number of RF-trees was set to 2,000. (a) Replicate analyses have been processed on ten reference tables. (b). The normalized mean absolute error (NMAE) is the absolute difference between the point estimate (here the median) and the (true) simulated value divided by the (true) simulated value.

**Figure S1.1.**
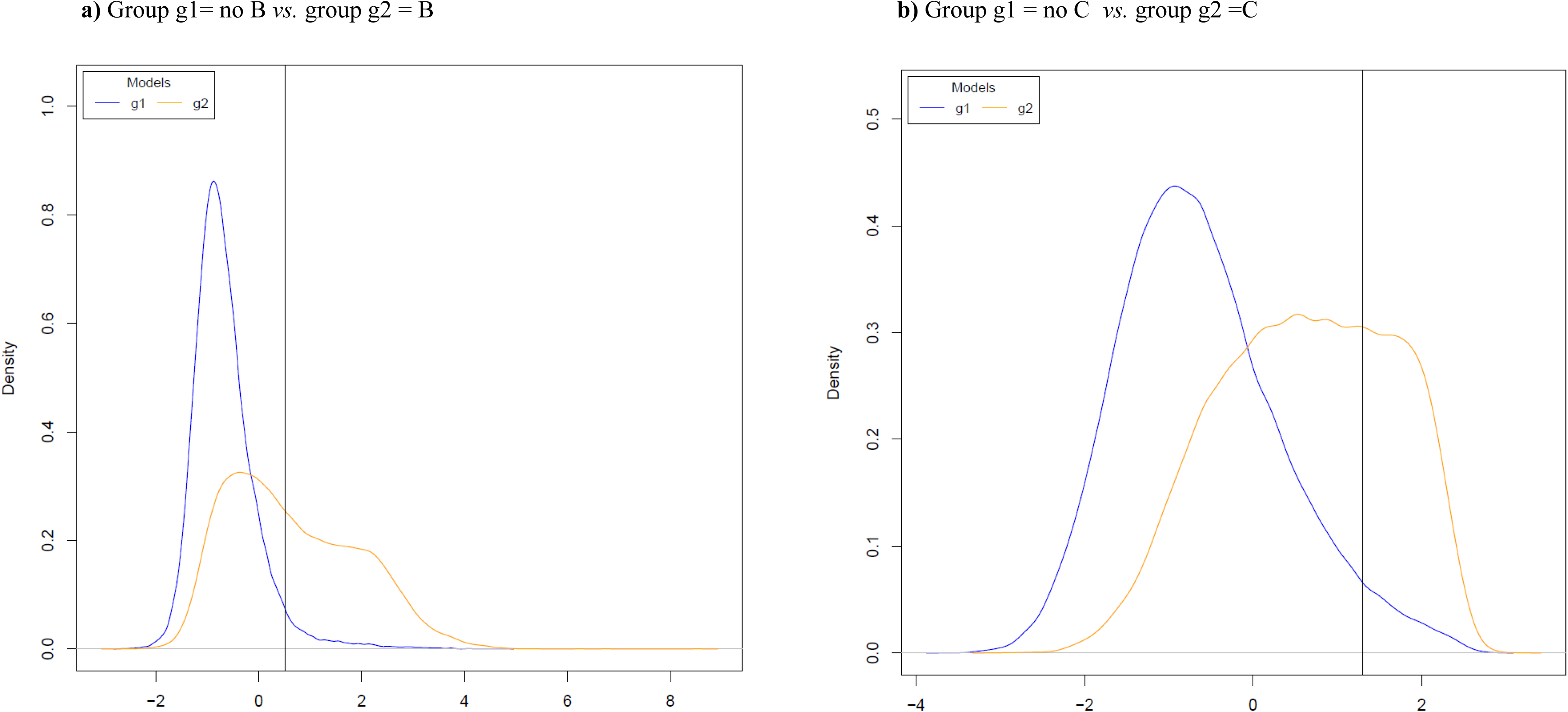
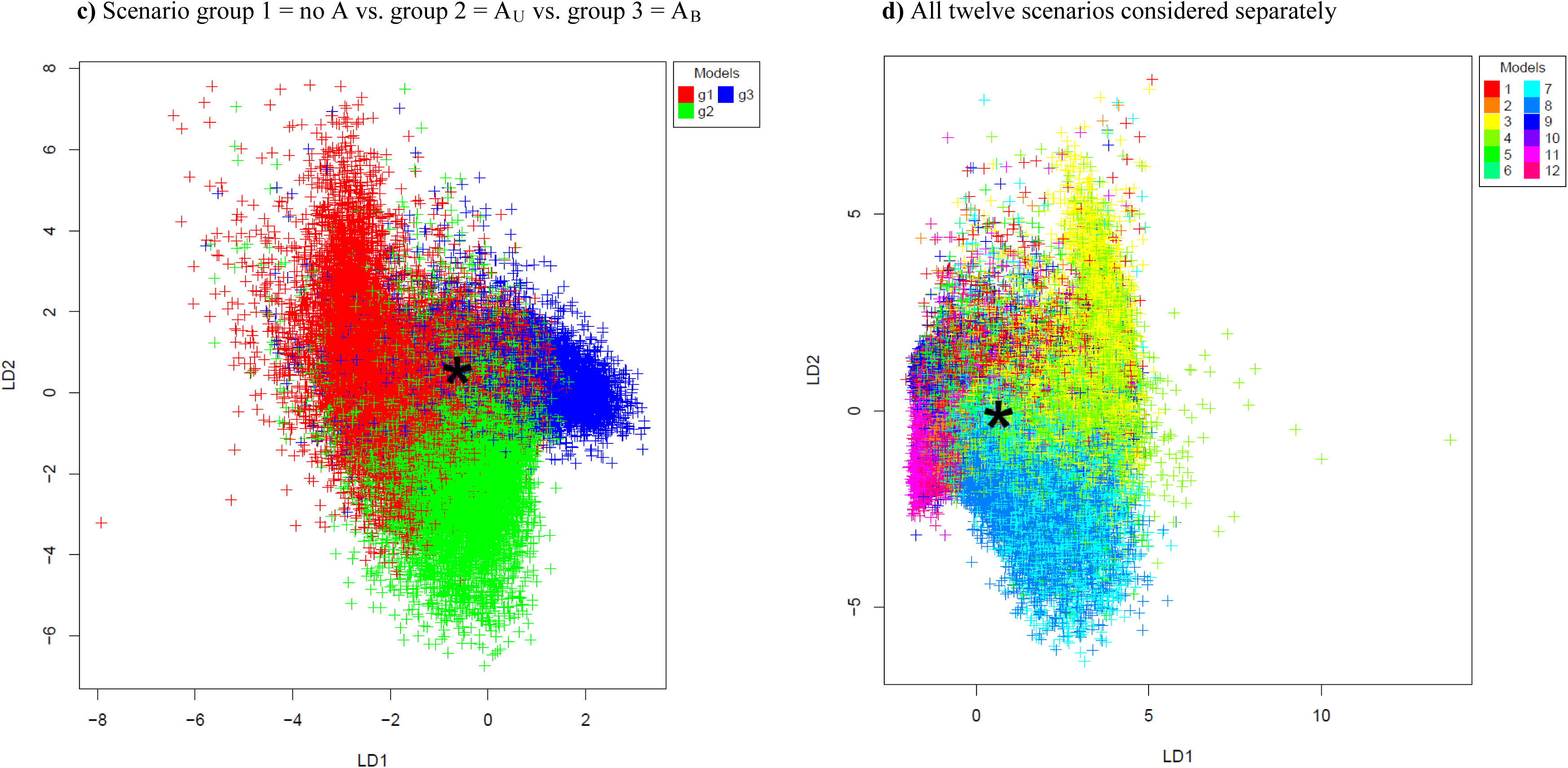
Projection on a single (when analyzing pairwise groups of scenarios) or on the first two LDA axes (when analyzing the twelve scenarios separately) of the observed dataset and the simulated datasets recorded in the reference table. Colors correspond to group of scenarios or individual scenarios. The location of the desert locust observed dataset is indicated by a vertical black line or a star. Scenarios were grouped based on the presence or not of a bottleneck in *S. g. flaviventris* (B or no B), a population size contraction in ancestor (C or no C) and a genetic admixture either unidirectional from *S. g. gregaria in* to *S. g. flaviventris* or bidirectional (A_U_ or A_B_ or no A). When considering the whole set of twelve scenarios separately d), the projected points substantially overlapped for at least some of the scenarios. This suggests an overall low power to discriminate among scenarios considered. Conversely, considering pairwise groups of scenarios, one can observe a weaker overlap of projected points (at least for a) and b)) suggesting a stronger power to discriminate among groups of scenarios of interest than when considering all scenarios separately. One can note that the location of the observed dataset (indicated by a vertical line) suggests an association with the scenario group with a bottleneck event in *S. g. flaviventris* and with the scenario group with a population size contraction in the ancestral population.

**Figure S1.2.**
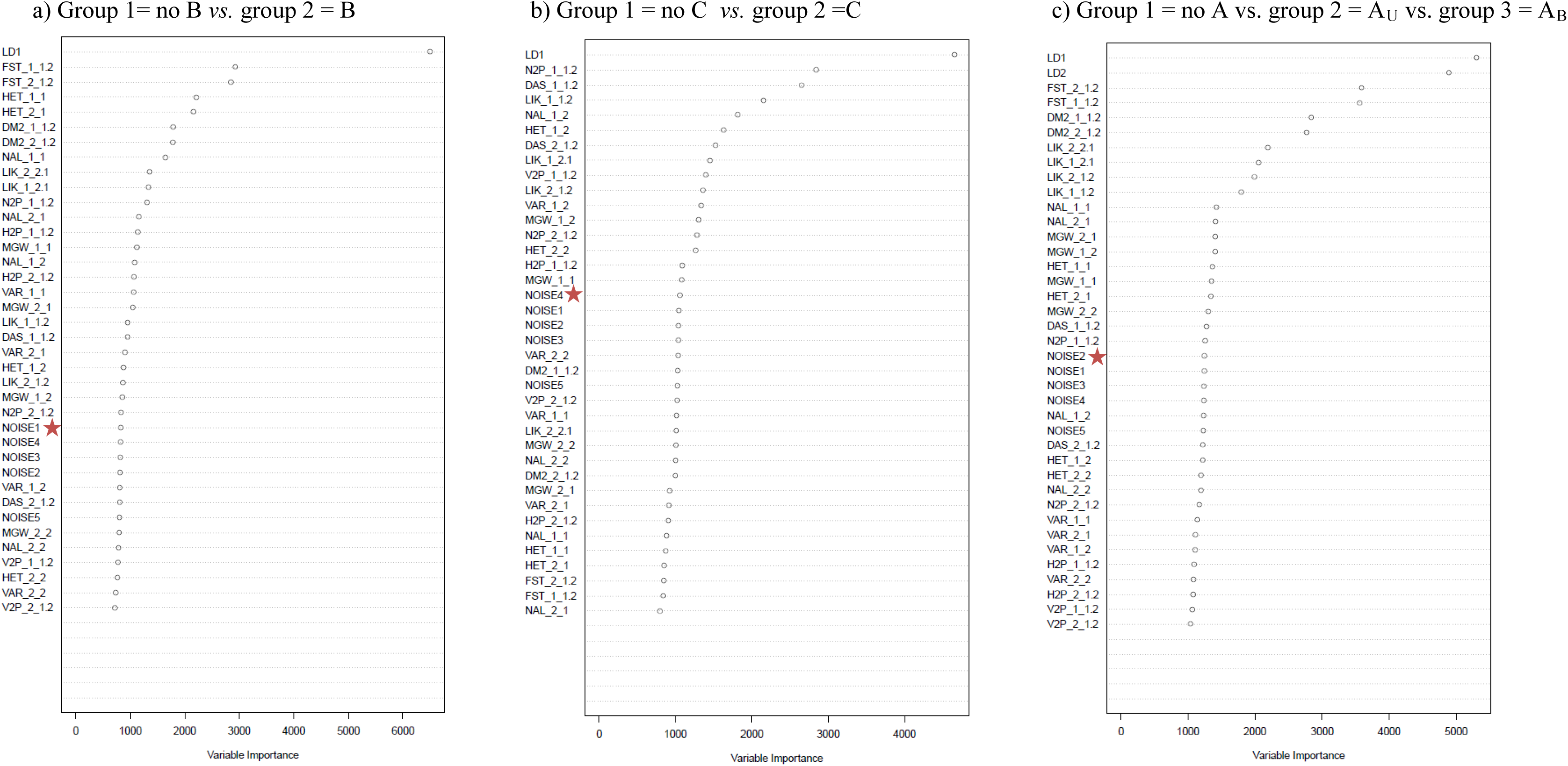
Contributions of ABC-RF summary statistics when choosing between groups of scenarios. The contribution of each 32 summary statistics and one LDA axis (or two LDA axes) is evaluated as the total amount of decrease in the Gini criterion (variable importance on the x-axis). The higher the contribution of the statistics, the more informative it is in the inferential process. The microsatellite set and subspecies sample are indicated at the end of each statistics by indices *k_i* for single population statistics and *k_i*.*j* for two population statistics, with *k*=1 for the set of untranscribed microsatellites or *k*=2 for the set of transcribed microsatellites, and *i*(*j*)=1 for the *S. g. flaviventris* subspecies or and *i*(*j*)=2 for the *S. g. gregaria* subspecies. See Table S1 for details on the summary statistics abbreviations. Five noise variables, randomly drawn into uniform distributions bounded between 0 and 1, and denoted NOISE1 to NOISE5 were added to the set of summary statistics processed by RF, in order to evaluate from which amount of decrease in the Gini criterion the summary statistics computed from our genetic datasets were not informative anymore (indicated by a red star). a) B = demographic bottleneck event, b) C = demographic contraction event and c) AU or AB or no A = unidirectional from *S. g. gregaria* into *S. g. flaviventris* or bidirectional genetic admixture event.

**Figure S1.3.**
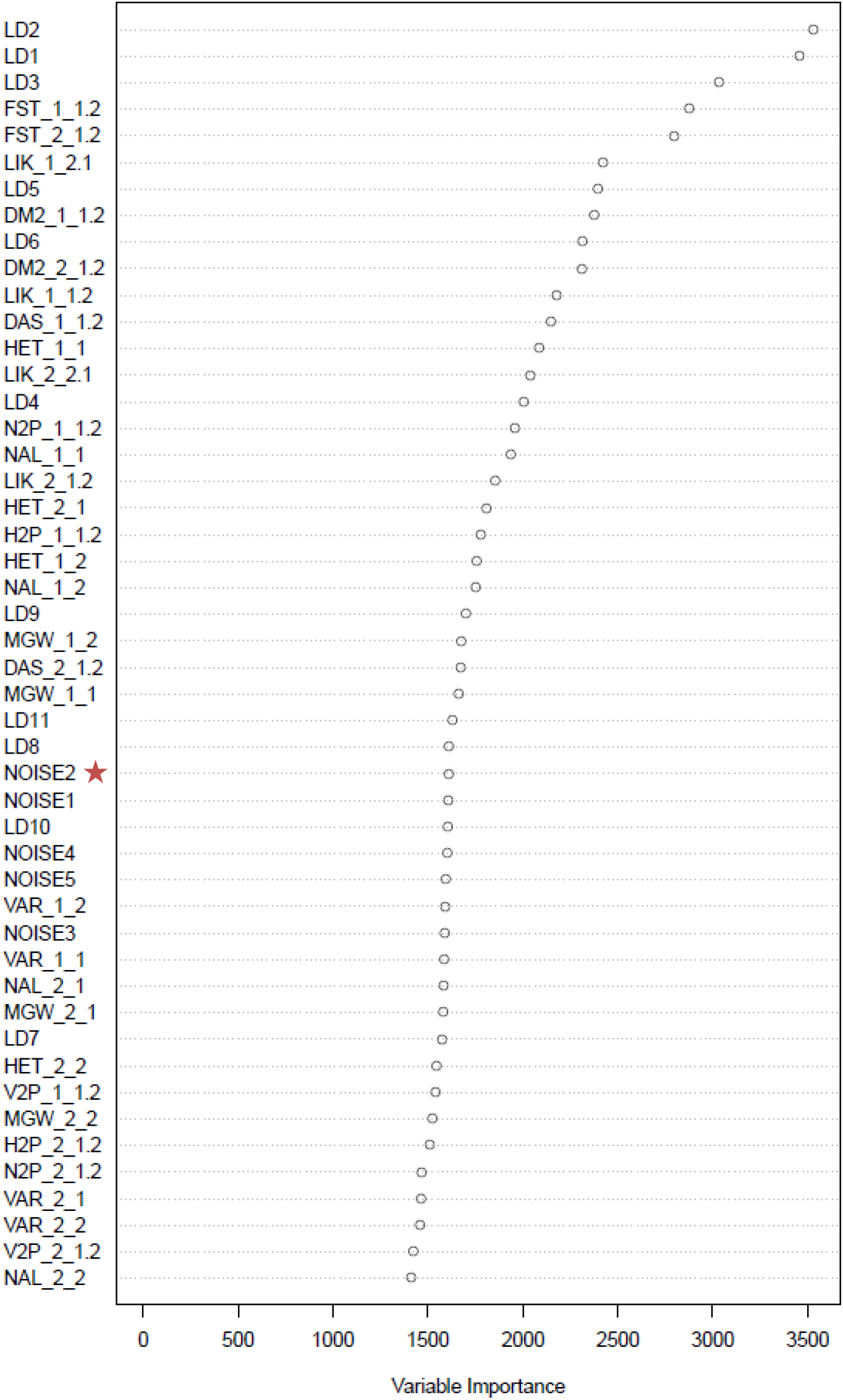
Contributions of ABC-RF summary statistics when choosing among the twelve individual scenarios. The contribution of each 32 summary statistics and eleven LDA axes is evaluated as the total amount of decrease in the Gini criterion (variable importance on the x-axis). The higher the contribution of the statistics, the more informative it is in the inferential process. The microsatellite set and subspecies sample are indicated at the end of each statistics by indices *k_i* for single population statistics and *k_i*.*j* for two population statistics, with *k*=1 for the set of untranscribed microsatellites or *k*=2 for the set of transcribed microsatellites, and *i*(*j*)=1 for the *S. g. flaviventris* subspecies or and *i*(*j*)=2 for the *S. g. gregaria* subspecies. See Table 1 for details on the summary statistics abbreviations. Five noise variables, randomly drawn into uniform distributions bounded between 0 and 1, and denoted NOISE1 to NOISE5 were added to the set of summary statistics processed by RF, in order to evaluate from which amount of decrease in the Gini criterion the summary statistics computed from our genetic datasets were not informative anymore (indicated by a red star).

**Figure S1.4.**
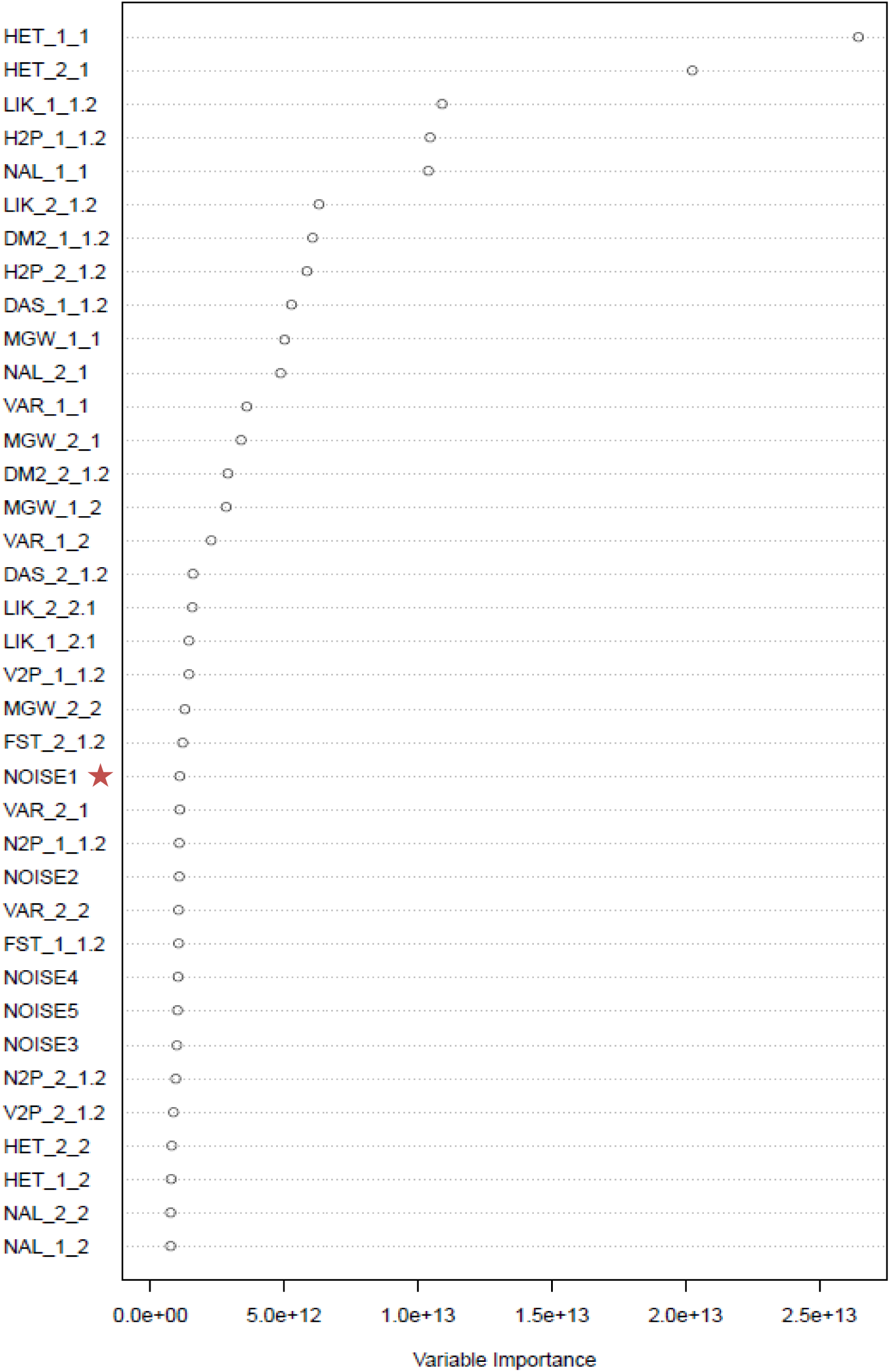
Contributions of ABC-RF summary statistics when estimating the divergence time between the two desert locust subspecies under the best supported scenario (scenario SCB). The contribution of each 32 summary statistics is evaluated as the total amount of decrease of the residual sum of squares, divided by the number of trees, (variable importance on the x-axis). The higher the contribution of the statistics, the more informative it is in the inferential process. The microsatellite set and subspecies sample are indicated at the end of each statistics by indices *k_i* for single population statistics and *k_i*.*j* for two population statistics, with *k*=1 for the set of untranscribed microsatellites or *k*=2 for the set of transcribed microsatellites, and *i*(*j*)=1 for the *S. g. flaviventris* subspecies or and *i*(*j*)=2 for the *S. g. gregaria* subspecies. See Table S6.1 for details on the summary statistics abbreviations. Five noise variables, randomly drawn into uniform distributions bounded between 0 and 1, and denoted NOISE1 to NOISE5 were added to the set of summary statistics processed by RF, in order to evaluate from which amount of decrease in the variable importance criterion the summary statistics computed from our genetic datasets were not informative anymore (indicated by a red star).

**Figure S1.5.**
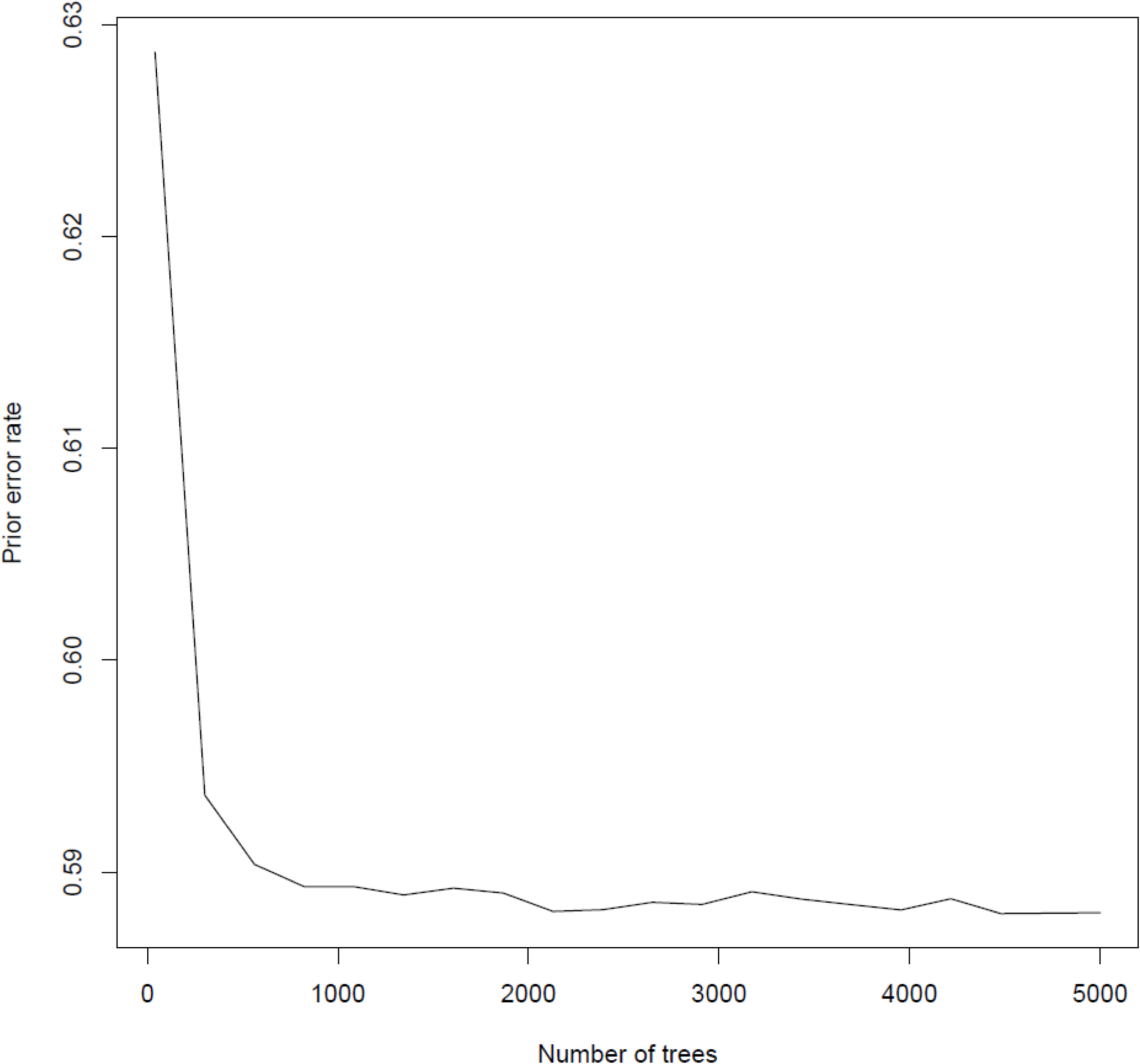
Effect of the number of RF-trees for scenario choice. We here illustrate the effect of the number of trees in the forest on the prior error rate when comparing the twelve scenarios separately. The number of datasets in the reference table simulated using DIYABC was 150,000. The shape of the curve shows that the prior error rate stabilizes for a number of RF-trees > 2,000.

**Figure S1.6.**
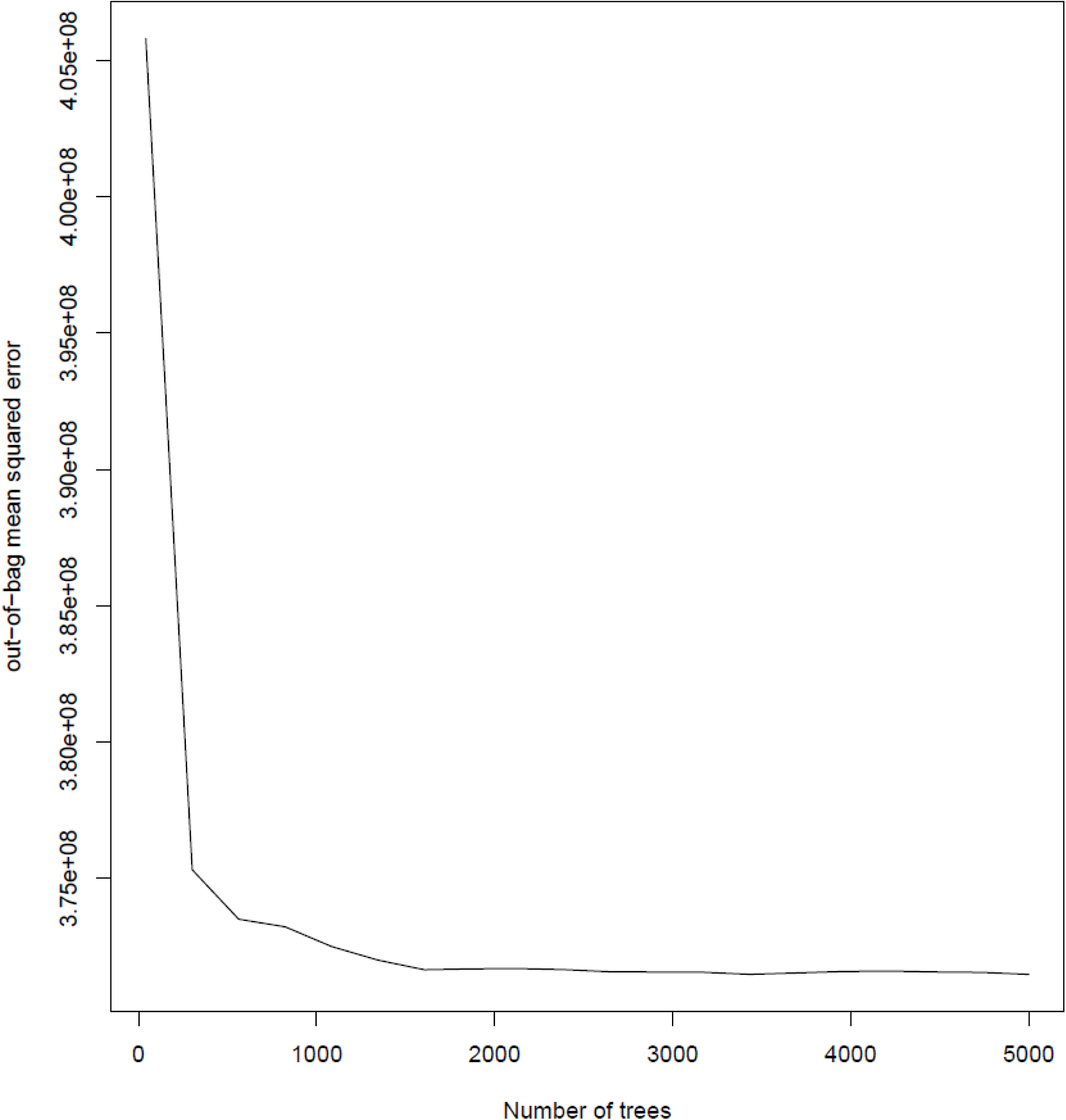
Effect of the number of RF-trees for parameter estimation. We here illustrate the effect of the number of trees in the forest on the out-of-bag mean square error of the divergence time between *S. g. gregaria* and *S. g. flaviventris* under the selected scenario SCB. The number of datasets in the reference table simulated using DIYABC was 100,000. The shape of the curve shows that the prior error rate stabilizes for a number of RF-trees > 1,500.

### Supplementary Material S2. Hindcasting the desert locust potential distribution into the mid-Holocene (HCO) and the last glacial maximum (LGM) using distribution modelling

#### Motivation

To complement distribution projections based on vegetation during the mid-Holocene (HCO; ∼6,000 years ago) and Last Glacial Maximum (LGM; ∼120,000 years ago) periods, we used species distribution modelling along with climatic reconstructions of the past. However, results from this modelling effort do not fully match vegetation reconstructions presented in the main text. Given that most Global Climate Models (GCMs) have been calibrated and tested largely with northern hemisphere data, uncertainties related to current and past extrapolations of climate over large areas of Africa are often unknown (*e*.*g*., Rowell *et al*. 2016). In this appendix, we present the results of the distribution modelling effort, while in the main text we base our formalization of evolutionary scenarios on paleo-vegetation reconstructions. Notice that the difference between the two approaches does not affect our major conclusions. Indeed, since paleo-vegetation maps show more severe changes of open vegetation distributions during glacial and interglacial periods, their use make us consider a wider spectrum of key biogeographic events than we would have identified based on climatic modelling (e.g. continuous colonization of southern Africa, population size contraction), and thus a larger set of possible scenarios. In addition, our inference of a most likely scenario including a long-distance migration event, and our estimation of a short time scale of divergence between the two clades, do not conflict with the results of any of these approaches (see also the section *On the influence of climatic cycles* in the Discussion of the main text).

#### Methods

In a previous study (Meynard *et al*. 2017), we used Climond data (Kriticos *et al*. 2012) to characterize the current distribution of the two desert locust subspecies, and project their potential fate under different climate change scenarios into 2070. Here, we could not use the same models because the climatic variables used to calibrate those models, in particular radiation and wetness indices, are not available into the mid-Holocene (HCO) and Last Glacial Maximum (LGM) periods. However, we apply here a similar modelling procedure with a different subset of environmental predictors. Instead of Climond, here we downloaded Worldclim v1.4 data (Hijmans *et al*. 2005) at a 5 min resolution (approximately 9 km at the equator).

For hindcasting, we first need to calibrate a model under current conditions, which can then be used to draw maps of potential ranges under HCO and LGM conditions. To do so, we used the occurrence data published in Meynard *et al*. (2017), which included decades of monitoring of *S. g. gregaria* in northern Africa, the middle east and southern Asia during remission periods, as well as field records and literature records of *S. g. flaviventris* in southern Africa. However, rather than modelling each clade independently as we did before, here we decided to group presence records for the two clades as a single unit for the following reasons: (1) here the emphasis is in paleo-climates, at a time scale where the two clades were not yet supposed to be differentiated; (2) a niche analysis in Meynard *et al*. (2017) did not show evidence of niche differentiation between *S. g. gregaria* and *S. g. flaviventris*, suggesting that the environment occupied by the southern clade is a subset of that occupied by the northern one; and (3) the biogeographic hypothesis supported by the molecular analysis presented here involves the colonization of the southern tip of south Africa by an isolated population of the northern sub-species, which is more generalist in terms of its environmental niche. We therefore considered that grouping occurrence records regardless of the subspecies would provide for a more accurate hindcasting scenario. However, preliminary analysis using separate modelling for each subspecies showed qualitatively similar projections when using *S. g. gregaria* alone (results not shown).

Finally, since here the emphasis is on Africa, we included only occurrence records in this continent, and used 10,000 pseudo-absences drawn at random from outside the combined current ranges of the two clades by using the distribution maps proposed in Meynard *et al*. (2017) to mask out areas available for pseudo-absence selection. We also assumed here that most of the African continent was accessible to the species during this geological time-scale, given the long known dispersal distances during outbreak periods, the wide distribution of the northern clade, and the fact that the species is present in both extremes of the continent. Consequently, here we do not delimit an accessible area, but we limit our modelling to continental Africa.

To model the species distribution, we used four climatic variables to represent mean and variability in temperature and precipitation, while minimizing correlation between them. The four variables selected using these criteria were annual mean temperature (BIO1), annual precipitation (BIO12), annual temperature range (BIO7) and precipitation seasonality (BIO15). We also tested the same forward variable selection procedure presented in Meynard *et al*. (2017), which provided very similar predictions but, for simplicity, we will not present those results here. Using these four climatic predictors, we applied three statistical models in Rv3.6.1 to draw a consensus map for all predictions: Generalized Additive Model (GAM) using library *mgcv*, and limiting the maximum model complexity to k=4 (Wood 2006); boosted regression trees (BRT) using the library *gbm*, with 2000 trees and no interactions (Elith *et al*. 2008); and MAXENT, using the library *dismo* and all default parameters *(Elith et al*. 2011). As mentioned above, background data was sampled at random in Africa from outside the species current distribution. Pseudo-absences received a combined weight of 50% when using GAM or BRT, in order to balance the large number of pseudo-absences with respect to presence records in the modelling process while representing the range of values of the environmental predictors in the study area (Barbet-Massin *et al*. 2012). The consensus was drawn as the median predicted probability between the three models, and four different threshold values were calculated from this consensus: the MST threshold (the threshold that maximizes the sum of sensitivity + specificity), often recommended as a good strategy to optimize presence and absence classification rates (Liu et al. 2005), was calculated to delimit the predicted range of the species and calculate classification rate statistics; and 3 different thresholds that represent the highest predicted values that include a fixed percentage of presences (50%, 75% and 90%). These thresholds were used to visualize on the maps the core (50% threshold) versus the more marginal (between 75% and 90% thresholds) habitats within the range. In this context, the 50% threshold represents the core distribution, i.e. the highest predicted values that contain more than half of the occurrences, the 90% threshold represents the area that includes most of the occurrences, and the area outside the 90% threshold represents marginal or no habitat, where the species is unlikely to occur. The area between the 75% and 90% threshold can be considered as habitat that is rather marginal but that is still part of the predicted range.

Classification success rates (AUC, TSS, sensitivity and specificity) were calculated for the consensus on all the dataset, using an MST threshold when needed. The thresholds calculated using current climate and occurrence data were then projected into past scenarios (HCO and LGM) using 3 different global circulation models at the same resolution: CCSM4, MIROC-ESM and MPI-ESM-P. We chose these three GCMs because their projections were available for both time steps of interest. Although the three GCMs are based on climatological principles, each one works in slightly different ways, and therefore produces different results. Consensus between GCMs presented below represents median values of predictions between the three GCMs.

#### Results

The current distribution of *S. gregaria*, as modelled using both subclades as a single unit and limited to continental Africa, is quite similar to the one already published in Meynard et al. (2017) (see Figure S2.1). There are, however, differences in the outline of the distributions, but the overall predicted area is largely overlapping and classification rates were excellent for this model under current conditions (% correctly classified = 87%; AUC = 0.960 ± 0.003; sensitivity = 0.911 ± 0.014; specificity = 0.870 ± 0.002; TSS = 0.788 ± 0.016). As expected, the species is more likely to occur in regions with mid to high temperatures, wide annual temperature range, low precipitation and high precipitation seasonality (Figure S2.2).

**Figure S2.1:**
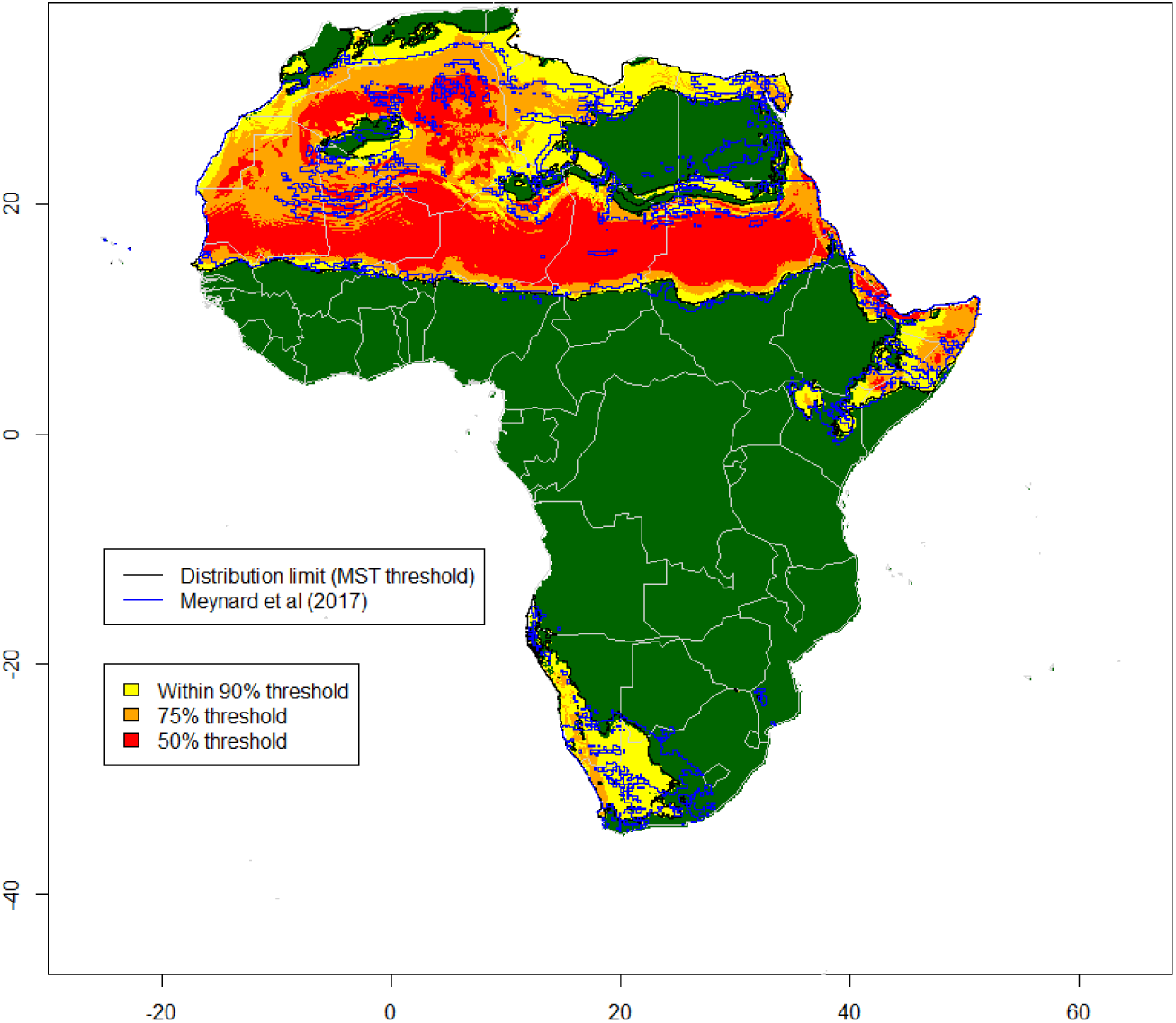
Current range as predicted by a model combining both subspecies’ occurrences and using four climatic predictors. The range predicted by this model (black contour) is compared to the one published in Meynard et al. (2017) using a different set of predictors and separating occurrences by subspecies (blue contour).

**Figure S2.2:**
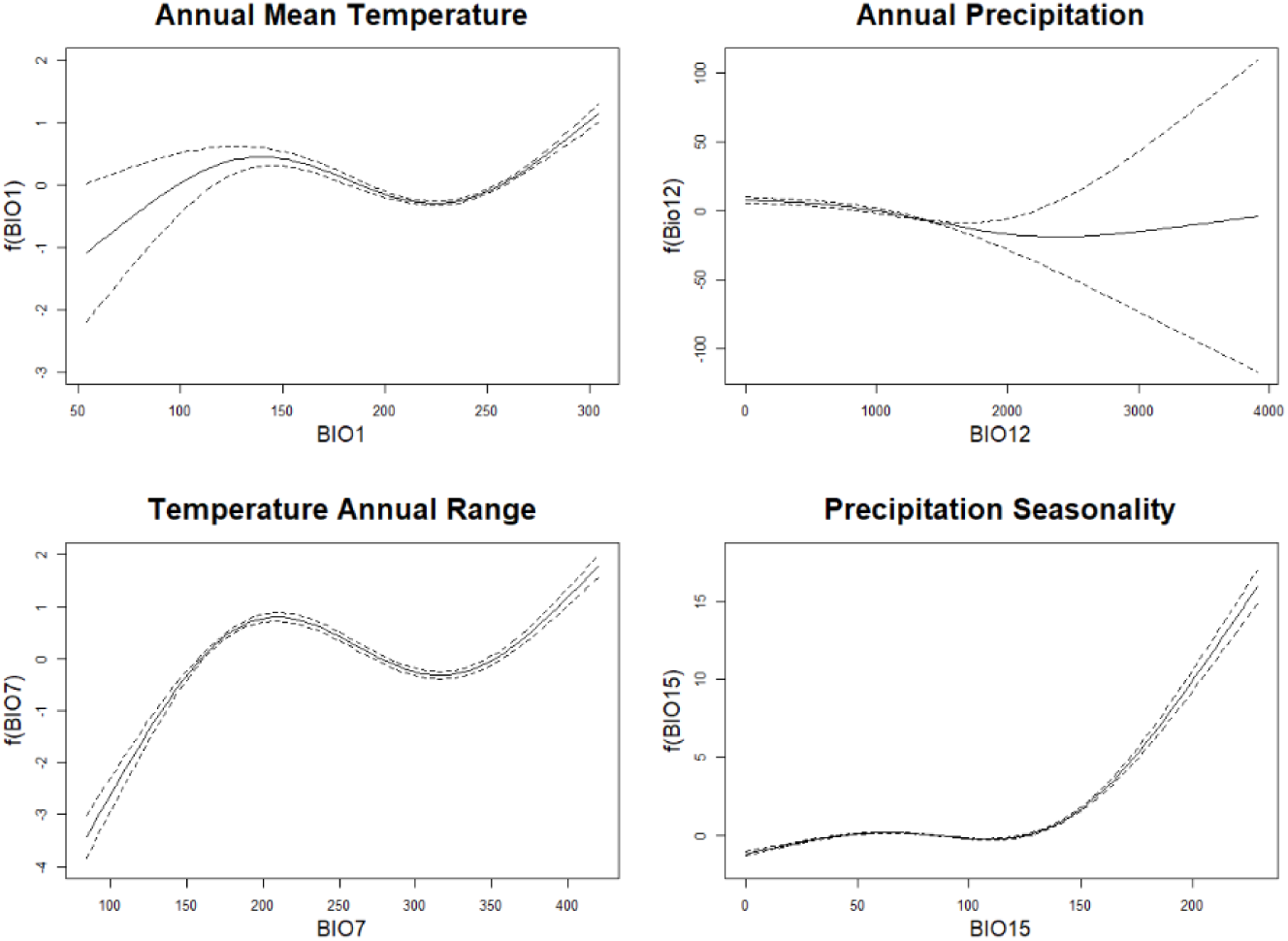
Response curves of S. gregaria against the four environmental predictors used in the modelling process, as derived from the GAM model. Notice that temperature is expressed as the real temperature x 10.

To better understand the results in terms of distributional changes, Figure S2.3 shows a comparison of the climatic variables used in the modelling process in the study region during the present, the HCO and the LGM periods across Africa. Although only one GCM scenario is shown (CCSM4), climatic conditions for the other two GCMs present similar trends. Notice that, although temperatures are thought to have been warmer than present during the HCO globally, those changes are not uniform across regions and are not reflected in these HCO scenarios for Africa. In all the scenarios in Figure S2.3, mean temperature showed a slight decrease across Africa, while precipitation increased slightly.

**Figure S2.3:**
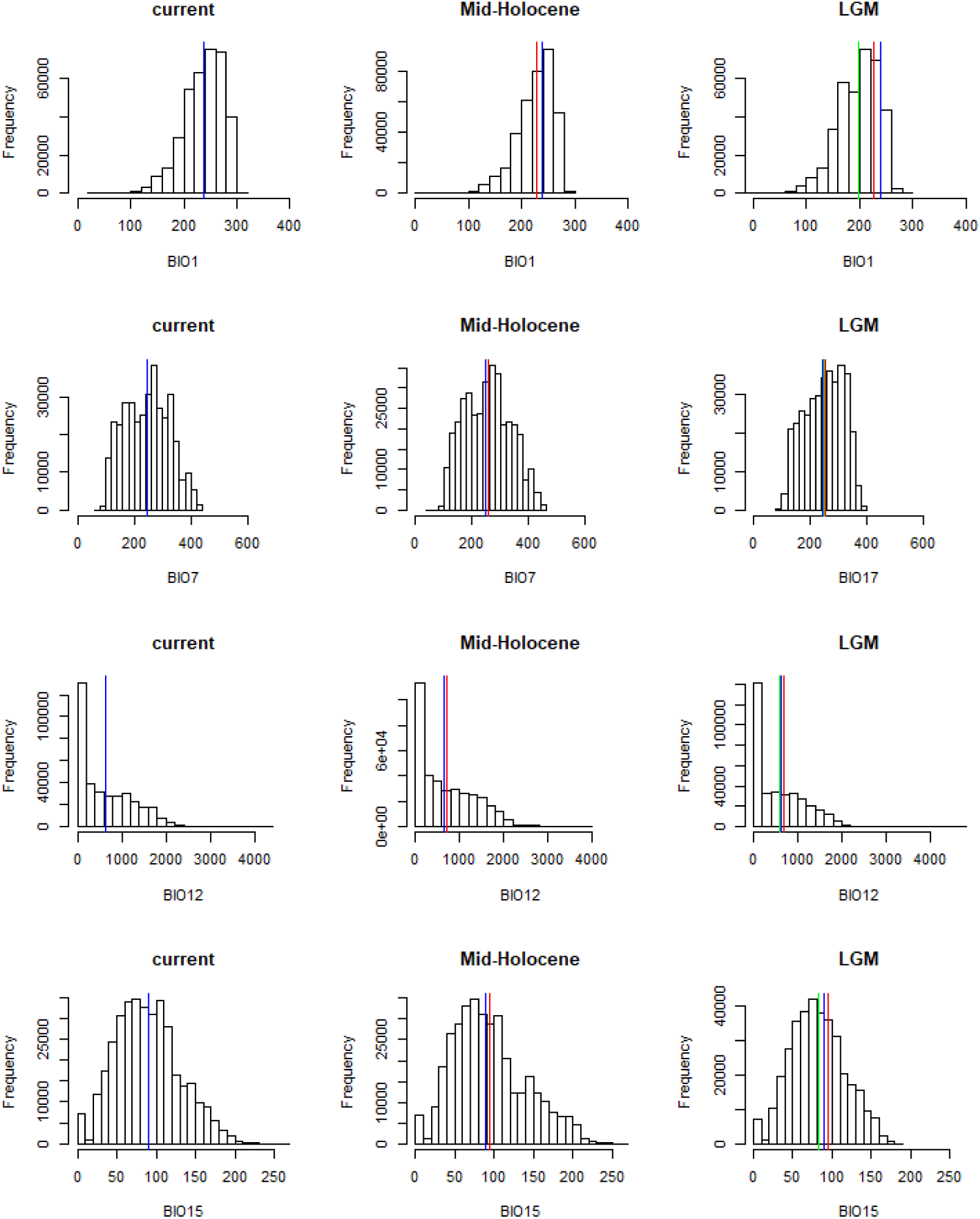
Comparison of climates across times (current climate, mid-Holocene, Last Glacial Maximum=LGM). Variables shown are those used in the modelling process. BIO1 = Mean Annual Temperature; BIO7 = Annual Temperature Range; BIO12 = Annual Precipitation; BIO15 = Precipitation Seasonality (Coefficient of Variation). Vertical color lines indicate mean values for each period: blue = current mean; red = HCO mean; green = LGM mean.

However, when looking at the distribution of those changes (Figure S2.4) some regions showed a decrease and others an increase in temperatures during the same period. Most of Africa shows changes that represent less than 10% of their current mean temperature values (Figure S2.4, upper left). Overall, conditions under HCO in Africa were therefore similar to current climatic conditions in terms of temperature but were moister in many regions that are relevant to the potential distribution of the desert locust (Figure S2.4, upper row).

**Figure S2.4:**
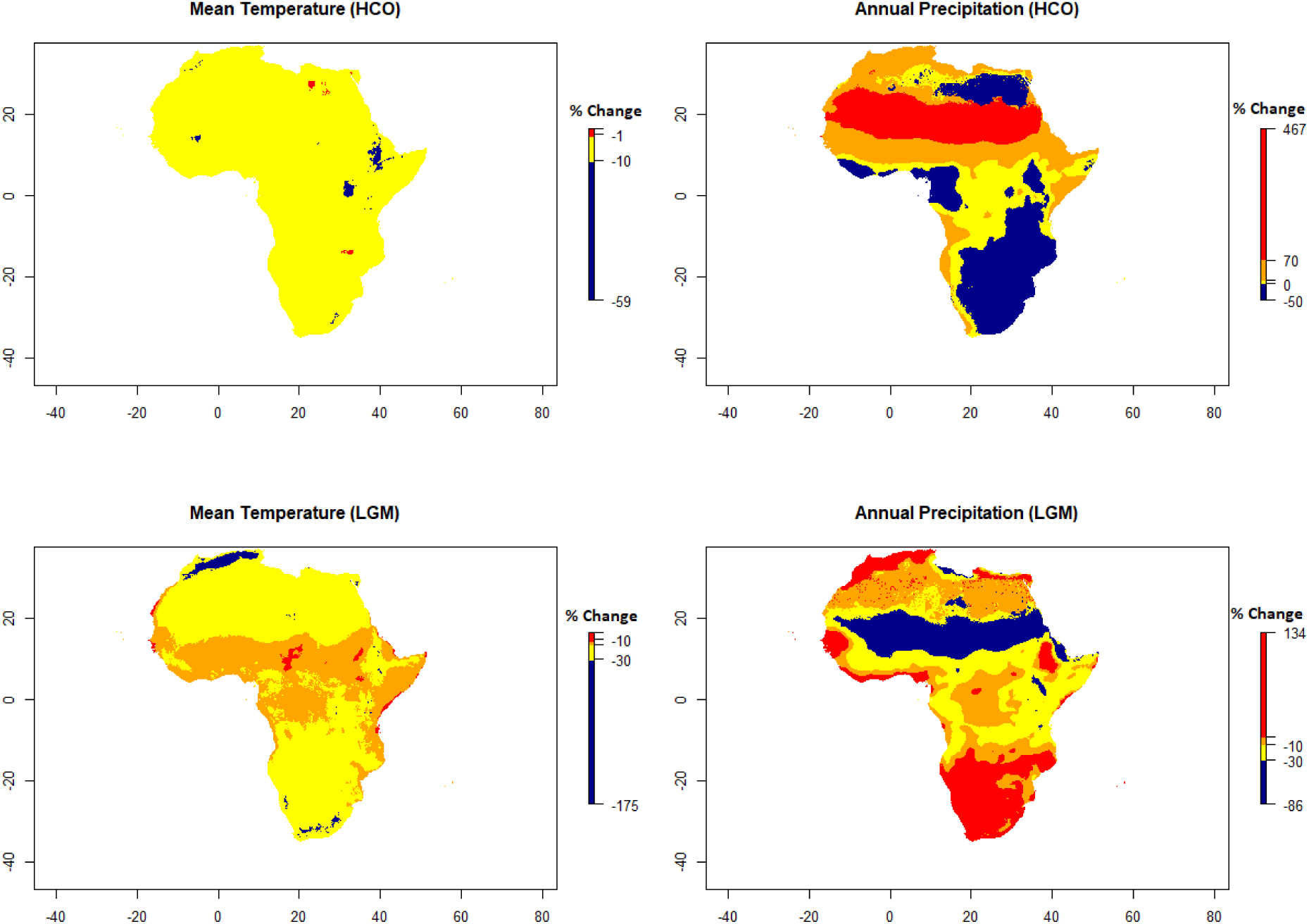
Relative changes in annual mean temperature (BIO1) and annual precipitation (BIO12) during the HCO and LGM, expressed as a percent of current values per grid cell. Positive values therefore indicate an increase whereas negative values represent a decrease with respect to current climatic conditions.

During the LGM (Figure S2.3, right column), mean temperature decreased more dramatically throughout all of Africa, while precipitation increased in some regions but decreased in others (Figure S2.4, lower row). While the entire continent experienced a decrease in temperature, changes in precipitation were more heterogeneous (Figure S2.4, lower right). Given that these changes are heterogeneous across regions, changes in the potential distribution of *S. gregaria* are less dramatic than one would expect given global simplifications of these changes.

Indeed, the predicted distribution during the HCO and the LGM are very similar to the current distribution (Figure S2.5). There is an overall shrinkage of the core habitat (in red in all maps), but the contour of the distribution (black contour) remains large and mostly overlapping with the current distribution.

**Figure S2.5:**
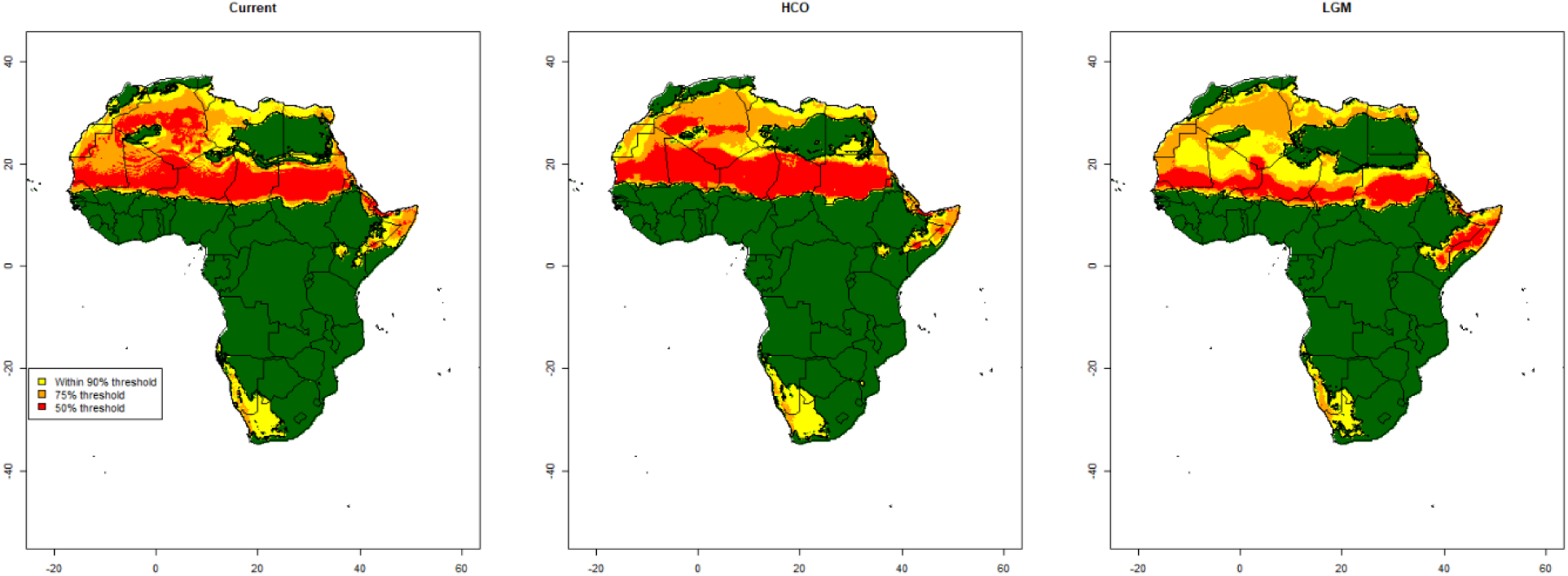
Comparison of current versus past potential distribution of S. gregaria during the HCO and LGM periods.

Although each GCM resulted in differences in terms of the projected distribution maps of *S. gregaria*, in the consensus distribution is projected to be very similar to the current distribution during the HCO, and slightly reduced during the LGM, with a shift of the core favorable climatic conditions towards a much reduced area, especially during the LGM (Figure S2.5).

Although climatic projections for each GCM result in different projected distributions during the HCO (Figure S2.6) and the LGM (Figure S2.7), overall the tendencies are the same: a similar distribution when comparing present and HCO conditions, and a more restricted potential distribution when considering LGM conditions.

**Figure S2.6:**
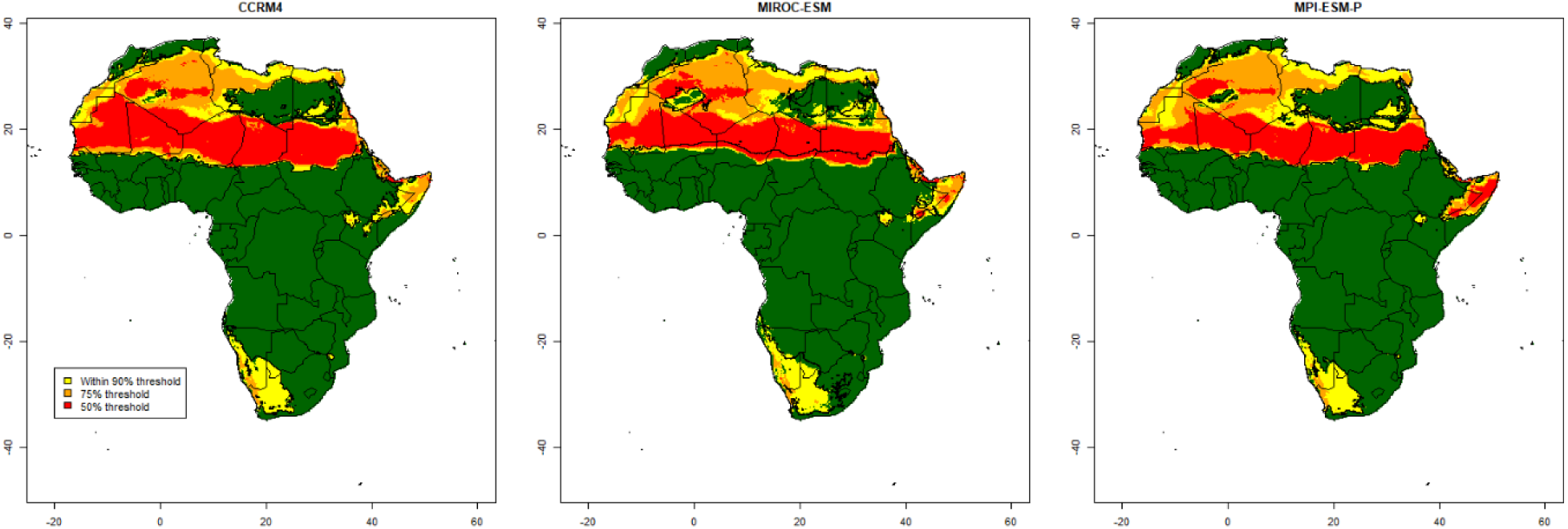
Projected potential distribution of S. gregaria during the mid-Holocene (∼6000 years BP) using three different Global Circulation Models (GCMs): CCRM4, MIROC-ESM and MPI-ESM-P.

**Figure S2.7:**
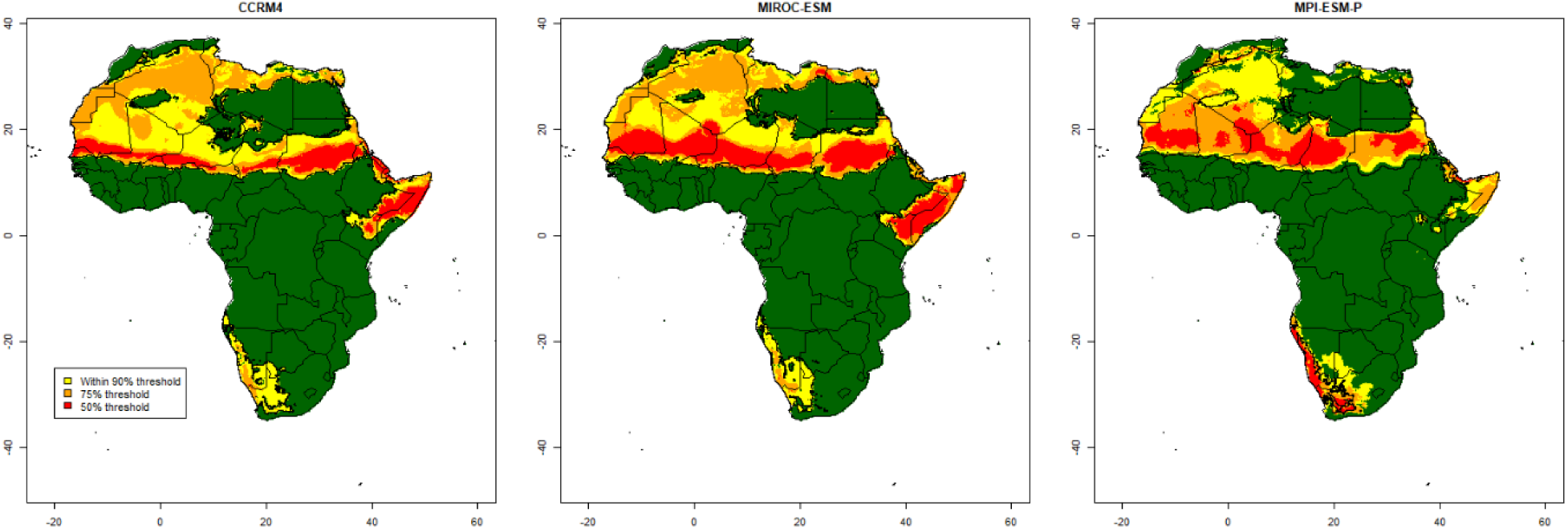
Projected potential distribution of S. gregaria during the LGM (∼20,000 years BP) using three different Global Circulation Models (GCMs): CCRM4, MIROC-ESM and MPI-ESM-P.

## Supplementary Material S3: Overview of the used ABC Random Forest (ABC-RF) methods

In this supplementary material, we provide readers with an overview of the Approximate Bayesian Computation Random Forest (hereafter ABC-RF) methods used in the present paper. We invite the reader to consult Pudlo et al. (2016), Estoup et al. (2018), and Raynal et al. (2019) for more in-depth explanations.

### ABC framework

Let **y** denote the observed data and ***θ*** a vector of parameters associated to a statistical model whose likelihood is *f* (. | ***θ***). Under the Bayesian parametric paradigm the posterior distribution

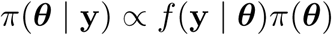

is of prime interest. It characterizes the distribution of ***θ*** given the observation **y** and can be interpreted as an update of the prior distribution *π*(***θ***) by the likelihood of **y**. The likelihood is hence pivotal, but unfortunately intractable in the evolutionary scenarios (models) we consider in the present study, as well as in many other evolutionary studies. As a matter of fact, the underlying Kingman’s coalescent process (Kingman, 1982) does not allow a close expression for the likelihood because all the possible genealogies and mutational process yielding **y** should be considered. To solve this issue, some likelihood-free methods have been developed using the fact that, even though the likelihood is not available, generating artificial (i.e. simulated) data for a given value of ***θ*** is much easier if not feasible (e.g. Beaumont (2010). Approximate Bayesian computation (ABC) is one of them (Beaumont et al., 2002).

In a nutshell, ABC consists in generating parameters ***θ***′ and associated pseudo-data **z** from the scenario, and accepting ***θ***′ as a realization from an *approximated* posterior if **z** is similar to **y**. In standard ABC treatments, the notion of similarity is defined through the use of a distance *ρ* to compare *η*(**z**) and *η*(**y**), where *η*(·) is a projection of the data in a lower dimensional space of summary statistics. Only pseudo-data providing distance lower than a threshold *ϵ* are retained. The choice of *ρ, η*(·) and *ϵ* is a major issue in ABC (Beaumont, 2010).

ABC-RF is a recently derived ABC approach based on the supervised machine learning tool named Random Forest (Breiman, 2001), which has as major advantage to avoid the three above-mentioned difficulties. Initially introduced in Pudlo et al. (2016) for model choice and then extended to parameter inference in Raynal et al. (2019), ABC-RF relies on the use of random forests on a set of simulated pseudo-data according to the generative Bayesian models under consideration. Let consider *M* Bayesian parametric models. For a given model index *m* ∈ {1, …, *M*}, a prior probability ℙ(ℳ = *m*) is defined, with ***θ***_*m*_ its associated parameters and *f*_*m*_(**y** | ***θ***_*m*_) its likelihood.

The generation process of a reference table made of *H* elements is described in Algorithm 1.

#### Algorithm 1

Generation of a reference table with *H* elements

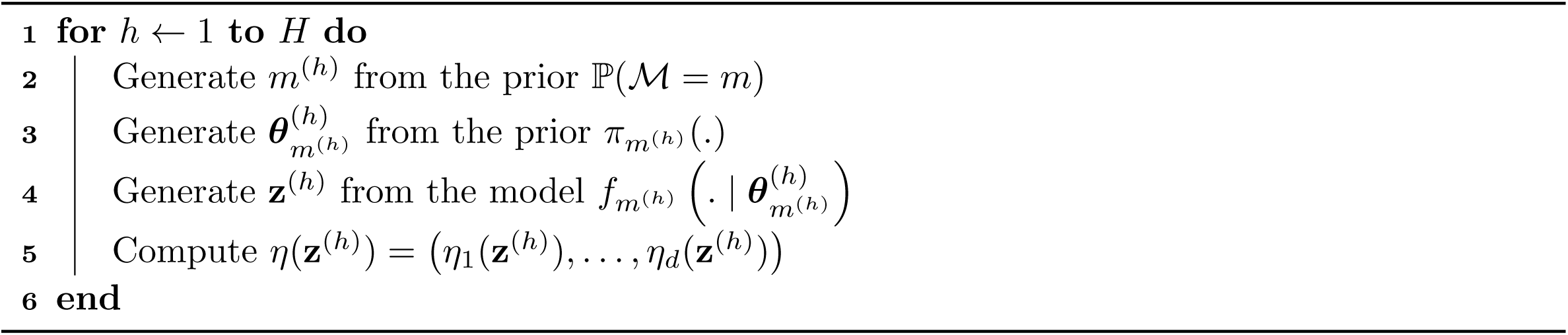

The output takes the form of a matrix containing simulated model indexes, parameters and summary statistics, as described below

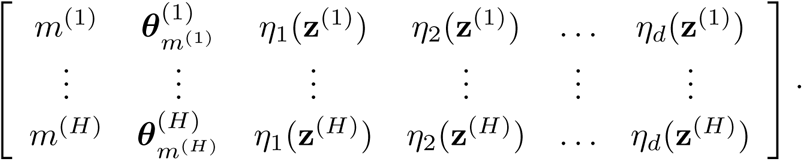

### ABC-RF for model choice

The ABC-RF strategy for model choice is described in Algorithm 2. The output is the affectation of **y** to a model (scenario), this decision being made based on the majority class of the RF tree votes.

#### Algorithm 2

ABC-RF for model choice

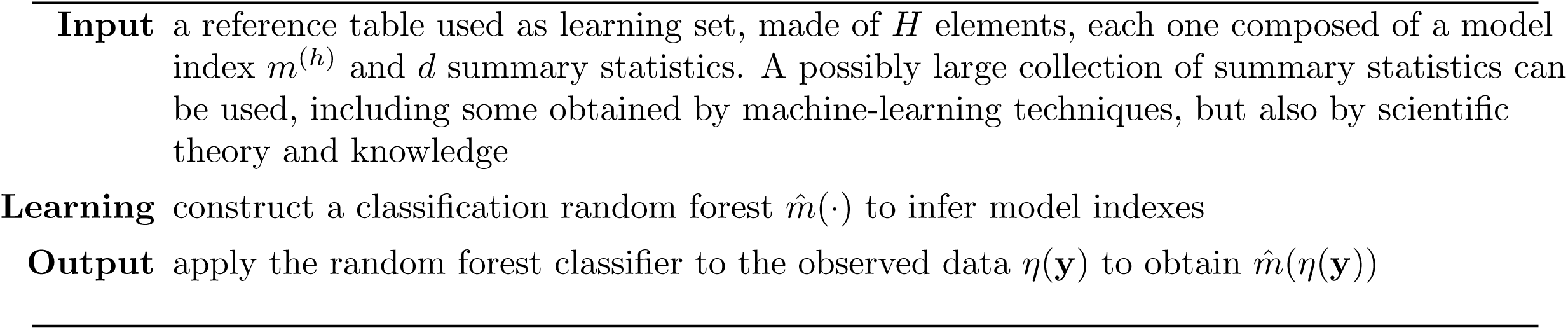

The selected scenario is the one with the highest number of votes in his favor. In addition to this majority vote, the posterior probability of the selected scenario can be computed as described in Algorithm 3.

Such posterior probability provides a confidence measure of the previous prediction at the point of interest *η*(**y**). It relies on the building of a regression random forest designed to explain the model prediction error. More specifically, and as a first step, posterior probability computation makes use of out-of-bag predictions of the training dataset. Because each tree of the random forest is built on a bootstrap sampling of the *H* elements of the reference table (i.e. the training dataset), there is about one third of the reference table that remains unused per tree, and this ensemble of left aside datasets corresponds to the out-of-bag. Thus, for each pseudo-data of the reference table, one can obtain an out-of-bag prediction by aggregating all the classification trees in which the pseudo-data was out-of-bag. In a second step, the out-of-bag predictions 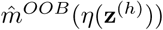 are used to compute the indicators 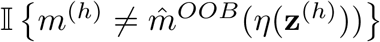. These 0 - 1 values are used as response variables for the regression random forest training, for which the explanatory variables are the summary statistics of the reference table. Predicting the observed data thanks to this forest allows the derivation of the posterior probability of the selected model (Algorithm 3). Note that using the out-of-bag procedure prevents over-fitting issues and is computationally parsimonious as it avoids the generation of a second reference table for the regression random forest training.

#### Algorithm 3

ABC-RF computation of the posterior probability of the selected scenario

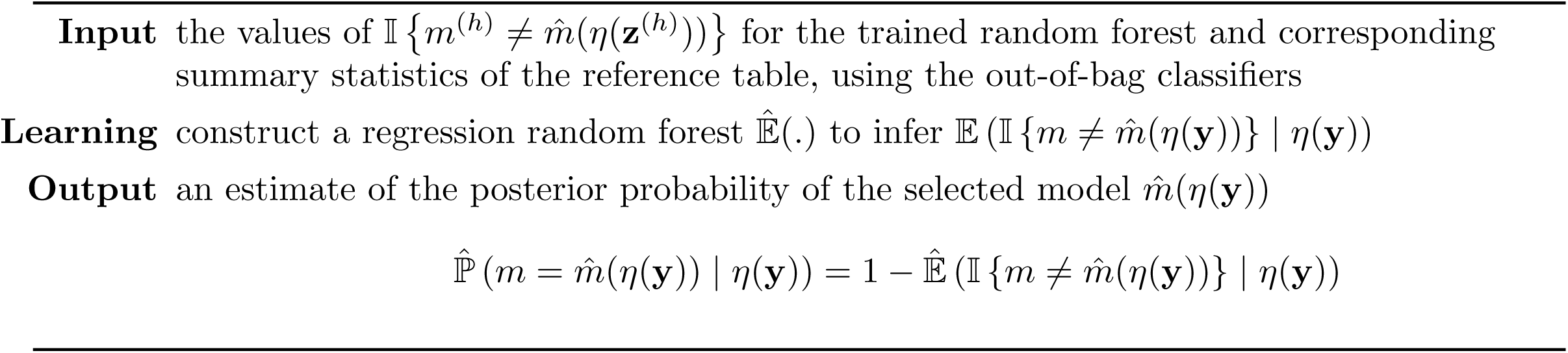

#### Model grouping

A recent useful add-on to ABC-RF has been the model-grouping approach developed in Estoup et al. (2018), where pre-defined groups of scenarios are analysed using Algorithm 2 and 3. The model indexes used in the training reference table are modified in a preliminary step to match the corresponding groups, which are then used during learning phase. When appropriate, unused scenarios are discarded from the reference table. This improvement is particularly useful when a high number of individual scenarios are considered and have been formalized through the absence or presence of some key evolutionary events (e.g. admixture, bottleneck, …). Such key evolutionary events allow defining and further considering groups of scenarios including or not such events. This grouping approach allows to evaluate the power of ABC-RF to make inferences about evolutionary event(s) of interest over the entire prior space and assess (and quantify) whether or not a particular evolutionary event is of prime importance to explain the observed dataset (see Estoup et al. (2018) for details and illustrations).

### ABC-RF for parameter estimation

Once the selected (i.e. best) scenario has been identified, the next step is the estimation of its parameters of interest under this scenario. The ABC-RF parameter estimation strategy is described in Algorithm 4 and takes a similar structure to Algorithm 2. The idea is to use a regression random forest for each dimension of the parameter space (i.e. for each parameter). For a given parameter of interest, the output of the algorithm is a vector of weights **w**_**y**_ that can be used to compute posterior quantities of interest such as expectation, variance and quantiles. **w**_**y**_ provides an empirical posterior distribution for *θ*_*m,k*_; see Raynal et al. (2019) for more details.

### Global prior errors

In both contexts, model choice or parameter estimation, a global quality of the predictor can be computed, which does not take the observed dataset (about which one wants to make inferences) into account. Random forests make it possible the computation of errors on the training reference table, using the out-of-bag predictions.

#### Algorithm 4

ABC-RF for parameter estimation

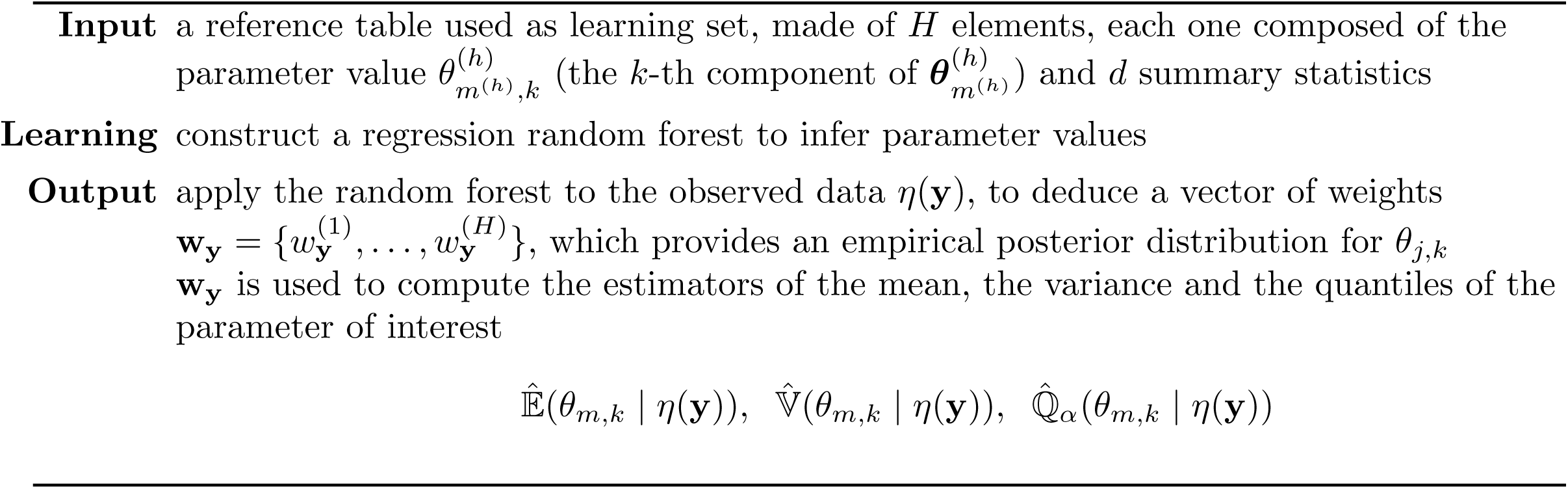

For model choice, this type of error is called the prior error rate, which is the mis-classification error rate computed over the entire multidimensional prior space. It can be computed as

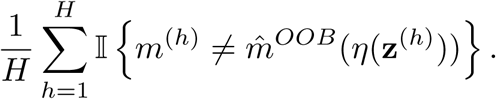

For parameter estimation, the equivalent is the prior mean squared error (MSE) or the normalised mean absolute error (NMAE), the latter being less sensitive to extreme values. These errors are computed as

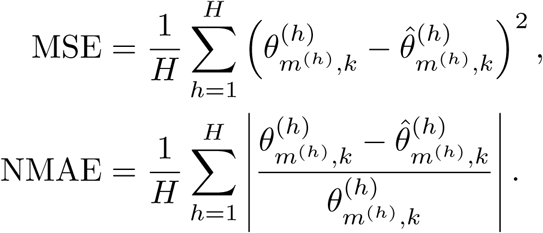

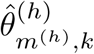 is the out-of-bag estimate of 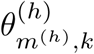. They can be perceived as Monte Carlo approximation of expectations with respect to the prior distribution.

### Local posterior errors

In the present paper, we propose some posterior versions of errors, which target the quality of prediction with respect to the posterior distribution. As such errors take the observed dataset *η*(**y**) into account, we mention them as local posterior errors.

For model choice, the posterior probability provided by Algorithm 3 is a confidence measure of the selected scenario given the observation. Therefore

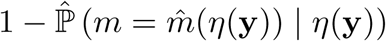

directly yields the posterior error associated to 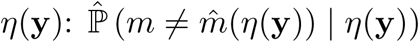.

For parameter estimation, when trying to infer on *θ*_*m,k*_, a point-wise analogous measure of a local error can be computed as the posterior expectations

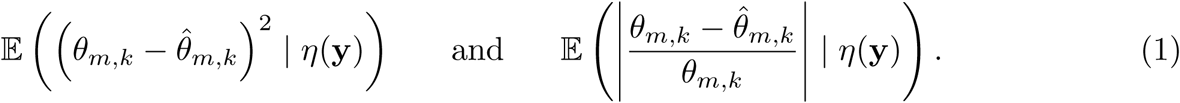

We approximate these expectations by

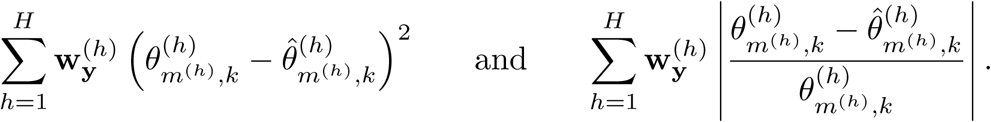

We again uses the out-of-bag information to compute 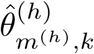, hence avoiding the (time consuming) production of a second reference table, and assume that the weights **w**_**y**_ from the regression random forest are good enough to approximate any posterior expectations of functions of *θ*_*m,k*_: 𝔼(*g*(*θ*_*m,k*_) | *η*(**y**)).

Another more expensive strategy to evaluate the posterior expectations (1) is to construct new regression random forests using the out-of-bag vector of values

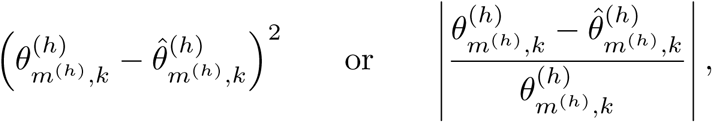

depending on the targeted error. The observation *η*(**y**) is then given to the forests, targeting the expectations (1).

Note that the values 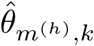 in the previous formulas can be replaced by either the approximated posterior expectations 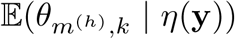 or the posterior medians 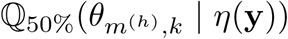, again using the out-of-bag information, to provide the local posterior errors. We found that both in the present paper (see main text, Materials and Methods section) and for various tests that we carried out on different inferential setups and datasets (results not shown), the posterior median provides a better accuracy of parameter estimation than the posterior expectation (aka posterior mean). This trends also holds for global prior errors that can be computed using either the mean or the median as point estimates.

As final comment, it is worth noting that so far a common practice consisted in evaluating the quality of prediction (for model choice or parameter estimation) in the neighborhood of the observed dataset, that is around *η*(**y**) and not exactly for *η*(**y**). For model choice, Estoup et al. (2018) use the so called posterior predictive error rate which is an error of this type. In this case, some simulated datasets of the reference table close to the observation are selected thanks to an Euclidean distance, then new pseudo-observed datasets are simulated using similar parameters, on which is computed the error (see also Lippens et al., 2017, for a similar approach in a standard ABC framework). However, the main problem of processing this way is the difficulty to specify the size of the area around the observation, especially when the number of summary statistics is large. We therefore do not recommend the use of such a “neighborhood” error anymore, but rather to compute the local posterior errors detailed above as the latter measured prediction quality exactly at the position of interest *η*(**y**).

## Supplementary Material S4. Details on results from ABC-RF treatments when using uniform priors for the three time period parameters of the studied scenarios

**Table S4.1.** Scenario choice for each of the ten replicate analyses using uniform priors for the three time period parameters of the studied scenarios.

| Reference table | Best scenario | Votes S | Votes SC | Votes SB | Votes SCB | Votes SA<sub>U</sub> | Votes SCA<sub>U</sub> | Votes SBA<sub>U</sub> | Votes SCBA<sub>U</sub> | Votes SA<sub>B</sub> | Votes SCA<sub>B</sub> | Votes SBA<sub>B</sub> | Votes SCBA<sub>B</sub> | Prior error rate | Posterior probability (best scenario) |
| --- | --- | --- | --- | --- | --- | --- | --- | --- | --- | --- | --- | --- | --- | --- | --- |
| 1 | scenario SCB | 0.013 | 0.045 | 0.147 | 0.462 | 0.007 | 0.022 | 0.020 | 0.040 | 0.041 | 0.099 | 0.034 | 0.069 | 0.778 | 0.596 |
| 2 | scenario SCB | 0.018 | 0.027 | 0.140 | 0.365 | 0.012 | 0.026 | 0.033 | 0.096 | 0.044 | 0.075 | 0.072 | 0.092 | 0.776 | 0.398 |
| 3 | scenario SCB | 0.012 | 0.077 | 0.182 | 0.326 | 0.006 | 0.030 | 0.019 | 0.069 | 0.038 | 0.102 | 0.064 | 0.074 | 0.775 | 0.426 |
| 4 | scenario SCB | 0.014 | 0.035 | 0.175 | 0.405 | 0.007 | 0.022 | 0.022 | 0.061 | 0.023 | 0.091 | 0.040 | 0.104 | 0.777 | 0.563 |
| 5 | scenario SCB | 0.019 | 0.048 | 0.101 | 0.416 | 0.010 | 0.024 | 0.026 | 0.063 | 0.039 | 0.092 | 0.055 | 0.106 | 0.778 | 0.547 |
| 6 | scenario SCB | 0.018 | 0.030 | 0.130 | 0.295 | 0.011 | 0.030 | 0.013 | 0.124 | 0.075 | 0.093 | 0.098 | 0.082 | 0.778 | 0.398 |
| 7 | scenario SCB | 0.016 | 0.057 | 0.094 | 0.308 | 0.011 | 0.018 | 0.015 | 0.050 | 0.028 | 0.113 | 0.058 | 0.231 | 0.775 | 0.511 |
| 8 | scenario SCB | 0.019 | 0.037 | 0.109 | 0.359 | 0.007 | 0.018 | 0.015 | 0.049 | 0.057 | 0.080 | 0.105 | 0.145 | 0.777 | 0.474 |
| 9 | scenario SCB | 0.010 | 0.052 | 0.114 | 0.286 | 0.017 | 0.023 | 0.014 | 0.070 | 0.069 | 0.136 | 0.098 | 0.113 | 0.777 | 0.460 |
| 10 | scenario SCB | 0.013 | 0.057 | 0.124 | 0.249 | 0.009 | 0.016 | 0.012 | 0.067 | 0.061 | 0.133 | 0.091 | 0.168 | 0.778 | 0.442 |
| **Mean** | **scenario SCB** | **0.015** | **0.047** | **0.132** | **0.347** | **0.010** | **0.023** | **0.019** | **0.069** | **0.048** | **0.101** | **0.072** | **0.118** | **0.777** | **0.481** |

**Table S4.2.**
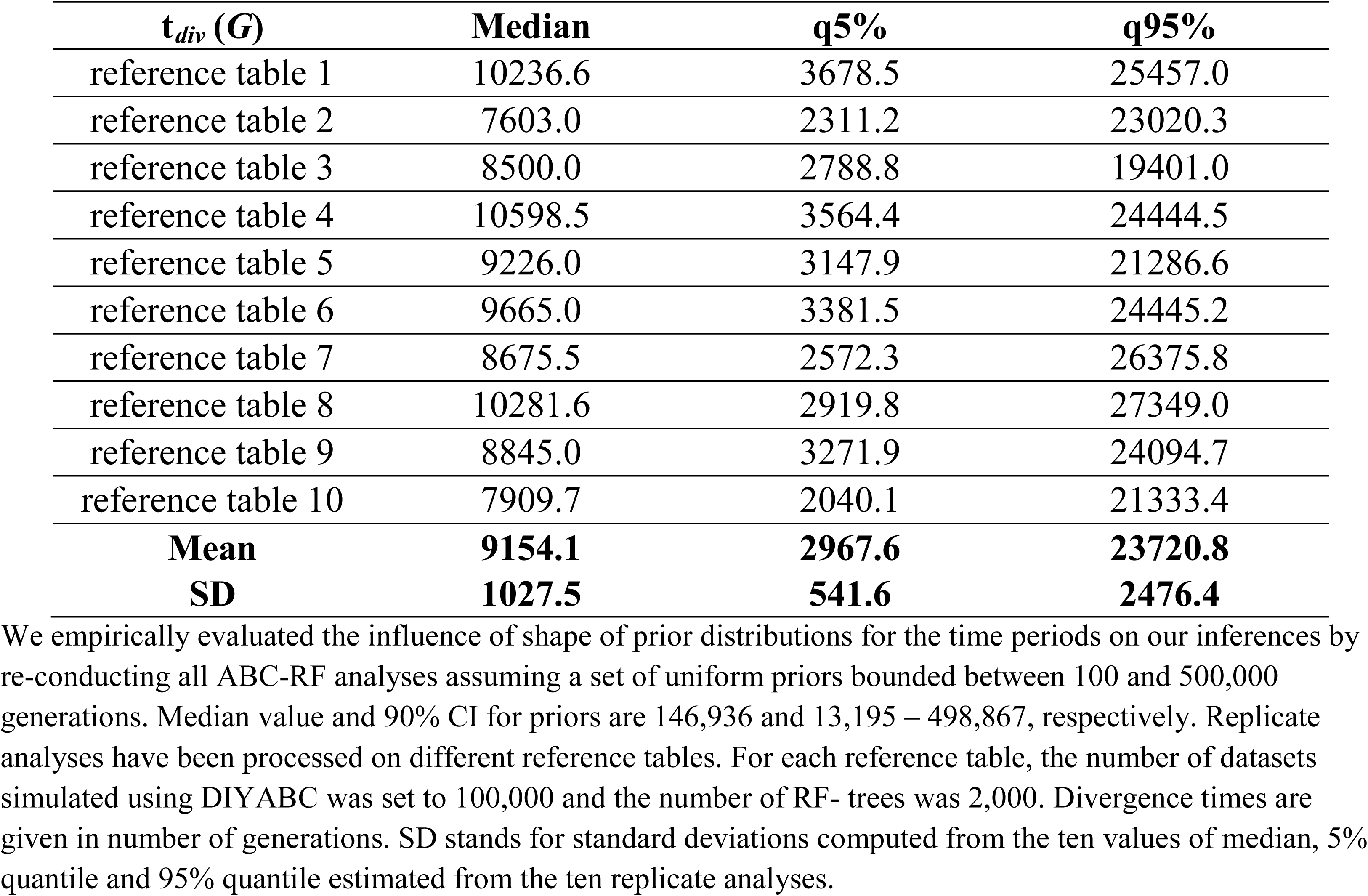
Estimation of the divergence time between *S. g. gregaria* and *S. g. flaviventris* for ten replicate analyses using uniform prior distributions for the three time period parameters under the best supported scenario (scenario SCB).

| $t_{div}$ (G) | Median | q5% | q95% |
| --- | --- | --- | --- |
| reference table 1 | 10236.6 | 3678.5 | 25457.0 |
| reference table 2 | 7603.0 | 2311.2 | 23020.3 |
| reference table 3 | 8500.0 | 2788.8 | 19401.0 |
| reference table 4 | 10598.5 | 3564.4 | 24444.5 |
| reference table 5 | 9226.0 | 3147.9 | 21286.6 |
| reference table 6 | 9665.0 | 3381.5 | 24445.2 |
| reference table 7 | 8675.5 | 2572.3 | 26375.8 |
| reference table 8 | 10281.6 | 2919.8 | 27349.0 |
| reference table 9 | 8845.0 | 3271.9 | 24094.7 |
| reference table 10 | 7909.7 | 2040.1 | 21333.4 |
| **Mean** | **9154.1** | **2967.6** | **23720.8** |
| **SD** | **1027.5** | **541.6** | **2476.4** |

We empirically evaluated the influence of shape of prior distributions for the time periods on our inferences by re-conducting all ABC-RF analyses assuming a set of uniform priors bounded between 100 and 500,000 generations. Median value and 90% CI for priors are 146,936 and 13,195 – 498,867, respectively. Replicate analyses have been processed on different reference tables. For each reference table, the number of datasets simulated using DIYABC was set to 100,000 and the number of RF- trees was 2,000. Divergence times are given in number of generations. SD stands for standard deviations computed from the ten values of median, 5% quantile and 95% quantile estimated from the ten replicate analyses.

**Figure S4.1.**
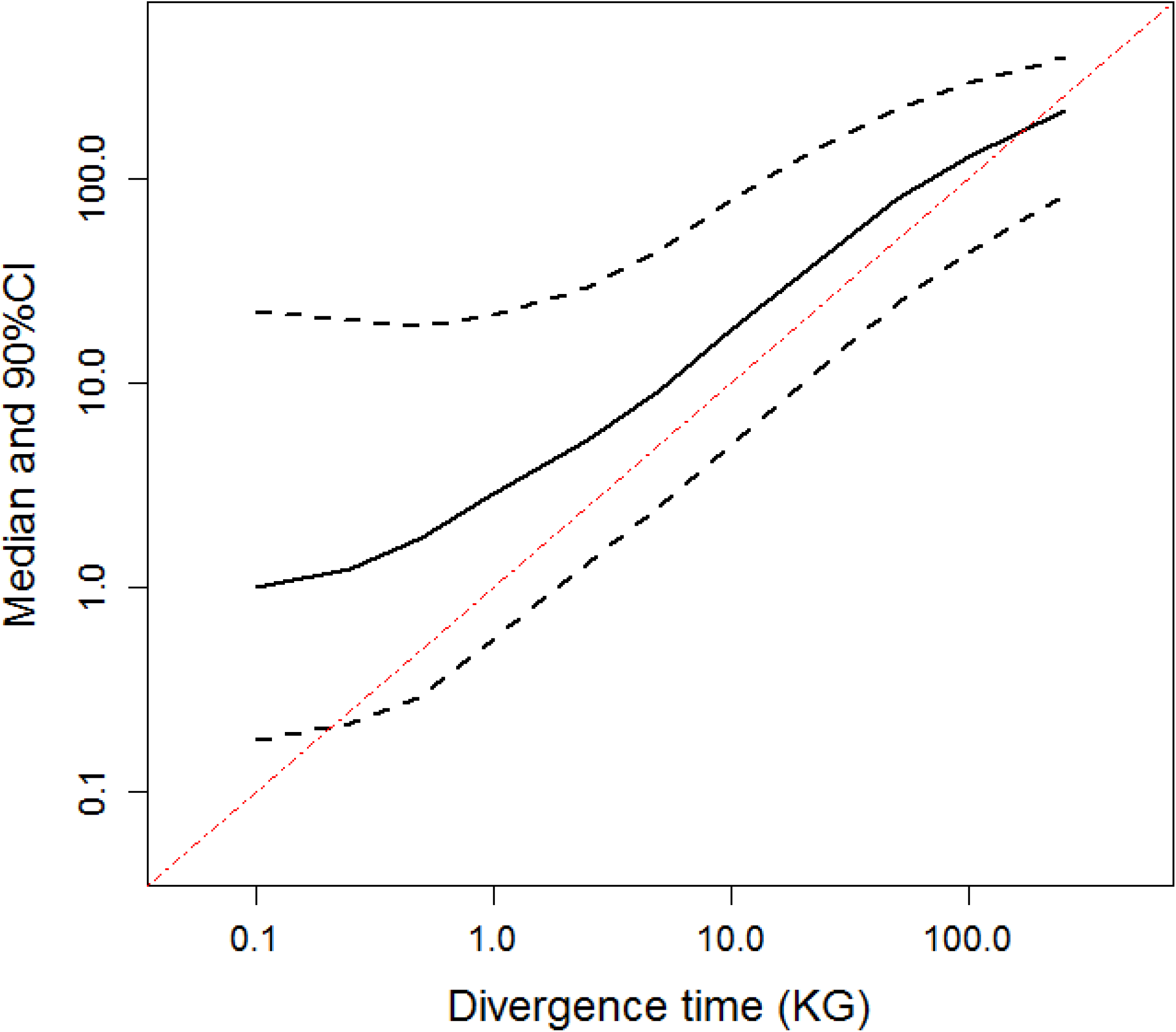
Estimation of the time since divergence between the two desert locust subspecies as a function of time scales using uniform prior distributions for the three time period parameters under the best supported scenario (scenario SCB). Simulated datasets (5,000 par divergence time) were generated for fixed divergence time values of 100; 250; 500; 1,000; 2,500; 5,000; 10,000; 25,000; 50,000; 100,000; and 250,000 generations. The median (plain lines) and 90% credibility interval (90% CI; dashed lines), averaged over the 5,000 datasets, are represented. Divergence time values are in number of generations.

## Notes

#### Summary of Updates

The Version 4 of this preprint has been peer-reviewed (by three reviewers), recommended by Peer Community In Evolutionary Biology (https://doi.org/10.24072/pci.evolbiol.100091), and formatted following the PCI Evol Biol template. Typos in some mathematical formula of the Supplementary Material S3 have been also corrected.

